# Biophysical characterization of the oligomeric states of recombinant Immunoglobulins type-M and their C1q binding kinetics by Biolayer Interferometry

**DOI:** 10.1101/2021.09.30.460443

**Authors:** Anne Chouquet, Andrea J. Pinto, Julia Hennicke, Wai Li Ling, Isabelle Bally, Linda Schwaigerlehner, Nicole M. Thielens, Renate Kunert, Jean-Baptiste Reiser

## Abstract

The Immunoglobulins type-M (IgMs) are one of the first antibody classes mobilized during immune responses against pathogens and tumor cells. Binding to specific target antigens enables the interaction with the C1 complex which strongly activates the classical complement pathway. This biological function is the basis for the huge therapeutic potential of IgMs but due to their high oligomeric complexity, *in vitro* production, biochemical and biophysical characterizations are challenging. In the present study, we present recombinant production of two IgM models (IgM617 and IgM012) in pentameric and hexameric states and the evaluation of their polymer distribution using different biophysical methods (AUC, SEC-MALLS, Mass Photometry and Transmission Electron Microscopy). Each IgM oligomer has individual specific expression pattern and yield with different protein quality likely due to intrinsic IgM properties and patterning. Nevertheless, the purified recombinant IgMs retain their ability to activate complement in a C1q dependent manner. And more importantly, a new method to evaluate their functional quality attribute by characterizing the kinetics of C1q binding to recombinant IgM has been developed using BioLayer Interferometry (BLI). We show that recombinant IgMs possess similar C1q binding properties as IgMs purified from human plasma.

## 1 Introduction

Since their discovery over a century ago, immunoglobulins (Igs) are considered to be one of the main components of the vertebrate humoral immune responses representing the first line of host protection against pathogens and altered-self cells, immune surveillance and homeostasis by recognizing and binding specific antigens. Immunoglobulins of type M (IgMs) are the first Igs produced and secreted by B cells in early infection and immune reaction stages in adults. They are highly oligomeric polypeptides composed of a heavy chain (H) and a light chain (L). Each chain contains an N-terminal variable domain and one (for the L chain) or four (for the H chain) C-terminal constant domains. IgMs are found as assembly of H_2_L_2_ protomer subunits either in a pentameric form (H_2_L_2_)_5_, containing an additional covalently linked peptide called the Joining (J) chain or, to a minor extent (less than 5%), in hexameric form (H_2_L_2_)_6_, devoid of J chain (for reviews Gong and Ruprecht, 2020; Jones et al., 2020; Keyt et al., 2020). IgM are also highly glycosylated with 5 to 6 different N-linked glycosylation chains that can represent up to 10 to 15% of their total molecular weight (Arnold et al., 2005). The overall assembly of IgMs is now revealed by modern high resolution Electron Microscopy (EM) methods: they have been observed as star-shaped structures with a compact Fc core and orbiting, labile and flexible Fab arms. While the hexamers appear to have the expected 6-fold symmetry, the pentamers possess a pseudo-hexameric symmetry where five IgM protomers occupy five of the six symmetric positions and the J chain the sixth position (Hiramoto et al., 2018; Li et al., 2020; Kumar et al., 2021).

In addition to their antigen recognition function, IgMs fulfill their immune effector function as one of the strongest activators of the classical pathway of the complement system. The binding of complexes between specific surface-exposed antigens and IgMs to the first recognition component of the classical pathway (CP), the C1q molecule, initiates complement-dependent cytotoxicity, a regulated amplifying proteolytic cascade which enables the membrane attack complex formation and the elimination of the pathogen or infected cell targets (Ricklin et al., 2010; Cedzyński et al., 2019). Found in high concentration in human serum, C1q, which forms the CP initiation C1 complex together with the pro-enzyme tetramer C1r_2_C1s_2_, is a 460kDa and highly flexible glycoprotein assembled from 18 polypeptide chains of 3 types (A, B, C). It is organized into six ABC heterotrimers forming six Ig recognition domains, the globular heads (gC1q), attached to collagen-like regions (CLR) allowing the assembly of what has been depicted as a “bouquet of tulips” or an “Eiffel-tower” (Kishore and Reid, 2000; Diebolder et al., 2014; Sharp et al., 2019).

Interaction studies of C1q with its Ig ligands led to the recent structures of Ig/C1 complexes by cryo-electron tomography and microscopy that finally revealed the general assembly of the CP initiation step and confirmed some, but not all, of the different biochemical studies performed to characterize the C1q/Ig binding properties and to identify the C1q/Ig binding sites (Ugurlar et al., 2018; Sharp et al., 2019). However, methods to measure the direct binding characteristics of C1q to Igs as functional quality attributes are rather limited in number (reviewed in Harboe et al., 2011). Traditional methods employ Enzyme-Linked Immunosorbent Assays (ELISA) to measure equilibrium and end-point deposition of Igs over coated C1q and vice-et-versa (exemplified in Duncan and Winter, 1988; Idusogie et al., 2000). More recently, label-free and real-time methods such as traditional Surface Plasmon Resonance (SPR) or novel BioLayer Interferometry (BLI) have been used to characterize the binding kinetics between C1q and IgGs (Moore et al., 2010; Patel et al., 2015; Zhou et al., 2018) or IgMs (Bally et al., 2019).

In the present study, we report (i) productions of our IgM models, IgM617 and IgM012 in pentameric and hexameric forms (ii) their biochemical and functional quality characterization using Analytical UltraCentrifugation (AUC), Size Exclusion Chromatography coupled to Multi-Angle Laser Light Scattering (SEC-MALLS), Mass Photometry (MP), Transmission Electron Microscopy (TEM) and in-house ELISA-like for detection of complement activation and (iii) the development of protocols to evaluate their bindings to C1q with BLI and to compare their kinetics characteristics.

## 2 Materials and Methods

### 2.1 IgM genetic constructs

IgM617 is originally expressed by the EBV-transformed B cell line HB617 and directed against glycosphingolipids overexpressed on the tumor cell surfaces and elicits cytotoxic T-cells (Vorauer-Uhl et al., 2010). IgM012 was developed after a class switch of a human IgG broadly-neutralizing HIV and is directed against the HIV1 envelope protein gp120 and in particular the carbohydrate mannose structure (Wolbank et al., 2003). All Light chains (L), Heavy chains (H) and Joining chain (J) cDNA of both IgM models were codon-optimized and sub-cloned into separate pIRES vectors as described in Chromikova *et al*. (Chromikova et al., 2015b), into pCEP4 vectors as described in Hennicke *et al*. (Hennicke et al., 2017) and into pcDNA3.1(+) vectors. For the last constructs, IgM chain cDNAs were amplified by PCR using conventional protocols and the pIRES constructs as templates. Additional flanked NheI and BamHI restrictions sites were also introduced by Polymerase Chain Reaction (PCR). The genes were then inserted individually with classical ligation techniques into pcDNA3.1(+) vectors (Invitrogen) containing a resistance cassette for either Geneticin (H chain), Zeocin (L chain) or Hygromycin (J chain).

### 2.2 Cell lines and expression of IgM models

IgM617-HLJ and IgM012-HLJ forms were expressed using either (i) CHO DG44 cell lines after co-transfection with pIRES constructs and generation of stable cells lines as described in Chromikova *et al*. (Chromikova et al., 2015b), (ii) HEK293E cell lines after co-transfection with pCEP4 vectors and transient expression as described in Hennicke *et al*. (Hennicke et al., 2019) or (iii) HEK293F after co-transfection or serial transfections with pcDNA3.1(+) vectors and generation of stable cell lines. Transfection and cultivation conditions of CHO DG44 and HEK293E cell lines have been described previously (Chromikova et al., 2015b; Hennicke et al., 2020). To generate stable IgM-expressing HEK293F cell lines, cells were grown in FreeStyle 293 expression medium and first co-transfected with H-chain- and L-chain-containing pcDNA3.1(+) plasmids using 293fectin according to the manufacturer’s protocol (Invitrogen). Stable transfectants producing either IgM617-HL or IgM012-HL were generated using cultivation in medium supplemented with 400 μg/ml G418 (Invitrogen) and 10 μg/ml Zeocin (Invitrogen). These cells were then transfected with J-chain-containing pcDNA3.1(+) plasmids in the same way and the stable transfectants producing either IgM617-HLJ or IgM012-HLJ were generated using cultivation in medium supplemented with additional 100 μg/ml hygromycin (Sigma-Aldrich). The stable cells expressing each IgM construct were then expanded in Freestyle 293 expression medium under antibiotic pressures every 3 to 4 days when cell density approached 3.10^6^ cells/ml.

### 2.3 Recombinant IgMs purification

All IgMs were purified from harvested supernatants according to Hennicke *et al*. (Hennicke et al., 2017). Briefly, POROS CaptureSelect™ IgM Affinity Matrix (Thermo Fisher Scientific) was used for affinity chromatography. Culture supernatants were directly applied to packed column and the IgMs were eluted with 1 M Arginine, 2 M MgCl_2_, pH 3.5 or pH 4.0. The collected fractions were immediately neutralized with 1M Tris pH 8.5. The IgM-containing fractions were then pooled and dialyzed against the next-step equilibration buffer. After concentration and for a second Size Exclusion Chromatography (SEC) step, they were applied on a Superose™ 6 increase 10/300 or 16/600 column (GE Healthcare/Cytiva) equilibrated in 0.1 M sodium phosphate pH 7.4, 0.2 M NaCl or in 0.025 M Tris-Base, 0.137 M NaCl, 0.003 M KCl, pH7.4 at a flow rate of 0.5 mL/min. Highly multimeric IgMs were eluted as a single peak separated from lower or higher oligomeric states and from nucleic acid contaminants. Identification of purified IgMs by SDS-PAGE was performed as described in Vorauer-Uhl *et al*. (Vorauer-Uhl et al., 2010) using Native PAGE 3-12% Bis-Tris gels and followed by Coomassie Blue staining.

### 2.4 C1q purification from plasma

C1q was purified from human serum according to the well-established protocol published by Arlaud *et al*. (Arlaud et al., 1979).

### 2.5 Analytical Ultracentrifugation

Sv-AUC experiments were conducted in an XLI analytical ultracentrifuge (Beckman, Palo Alto, CA) using an ANTi-60 rotor and double channel Ti centre pieces (Nanolytics, Germany) of 12- or 3-mm optical path length equipped with sapphire windows, the reference channel being typically filled with the sample solvent. Acquisitions were done overnight at 4°C and at 20000 rpm (32000 g) using absorbance (280 nm) and interference detection. Data processing and analysis were done using the program SEDFIT (Schuck, 2000) from P. Schuck (NIH, USA), REDATE (Zhao et al., 2015) and GUSSI (Brautigam, 2015) from C. Brautigam (USA), and using standard equations and protocols described previously (Salvay et al., 2008; Le Roy et al., 2013, 2015).

### 2.6 Size Exclusion Chromatography - Multi Angle Laser Light scattering (SEC-MALLS) analyses

SEC combined with online detection by MALLS, refractometry and UV-Vis was used to measure the absolute molecular mass in solution. The SEC runs were performed using a Superose™ 6 increase 10/300 column (GE Healthcare/Cytiva) equilibrated in 0.025 M Tris-Base, 0.137 M NaCl, 0.003 M KCl, pH 7.4. Separation was performed at room temperature and 50 μl of protein sample, concentrated to about 1 mg/ml, were injected with a constant flow rate of 0.5 ml/min. Online MALLS and differential refractive index detection were performed with respectively a DAWN-HELEOS II detector (Wyatt TechnologyCorp.) using a laser emitting at 690 nm and an Optilab T-rEX detector (Wyatt Technology Corp.), respectively. Weight-averaged molar masses (Mw) determination was done with the ASTRA6, using the “protein conjugate” module of the ASTRA software package. The following refractive index increments and UV-Vis absorbance values were used: *dn/dc* protein = 0.185 mL/g; *dn/dc* glycosylation = 0.15 mL/g; A_280_=1.38 ml/mg.cm.

### 2.7 Mass Photometry

Coverslips (high precision glass coverslips, 24×50 mm^2^, No. 1.5H; Marienfeld, Lauda-Königshofen, Germany) were cleaned by sequential sonication in Milli-Q H_2_O, 50% iso-propanol (HPLC grade)/Milli-Q H_2_O, and Milli-Q H_2_O (5 min each), followed by drying with a clean nitrogen stream. To keep the sample droplet in shape, reusable self-adhesive silicone culture wells (Grace Bio-Labs reusable CultureWell™ gaskets) were cut in segments of 4 to 10. To ensure proper adhesion to the coverslips, the gaskets were dried well by using a clean nitrogen stream. To prepare a sample carrier, gaskets were placed in the center of the cleaned coverslip and fixed tightly by applying light pressure with the back of a pipette tip. Protein landing was recorded using a Refeyn One^MP^ (Refeyn Ltd., Oxford, UK) MP system by forming a droplet of each IgM sample with a final concentration of 10 nM in a buffer solution containing 0.025 M Tris-Base, 0.137 M NaCl, 0.003 M KCl, pH7.4. Movies were acquired for 120 s (12000 frames) with Acquire^MP^ (Refeyn Ltd., v2.1.1) software using standard settings. Contrast-to-mass (C2M) calibration was performed using a mix of proteins with molecular weight of 66, 146, 500, and 1046 kDa. Data were analyzed using Discover^MP^ (Refeyn LTD, v2.1.1) and analysis parameters were set to T1 = 1.2 for threshold 1. The values for number of binned frames (nf = 8), threshold 2 (T2 = 0.25), and median filter kernel (=15) remained constant. The mean peak contrasts were determined in the software using Gaussian fitting. The mean contrast values were then plotted and fitted to a line. The experimental masses were finally obtained by averaging replicates and errors were the standard deviation.

### 2.8 Transmission electron microscopy (TEM)

About 4 μl of diluted IgM samples (60 to 80 ng) were applied to a carbon film evaporated onto a mica sheet. The carbon film was then floated off the mica in ∼100 μL 2 % sodium silicotungstate (SST, Agar Scientific) and transferred onto a 400 mesh Cu TEM grid (Delta Microscopies). Images were acquired with a CETA camera on a Tecnai F20 TEM microscope operating at 200 keV.

### 2.9 Complement activation-ELISA

The activation of the classical complement pathway was monitored by an ELISA based on the detection of C4b deposition according to Bally *et al*. and Hennicke *et al*.(Bally et al., 2019; Hennicke et al., 2020). Briefly, 200 ng of IgMs were immobilized on a MaxiSorp 96-well plate (Thermo Fisher Scientific) and incubated with Normal Human Serum (NHS) diluted 25 times, C1q-depleted serum (NHSΔ, CompTech) diluted 25 times or NHSΔ diluted 25 times reconstituted with purified human C1q (4 μg/mL). NHS was obtained from the Etablissement Français du Sang Rhône-Alpes (agreement number 14-1940 regarding its use in research). Unspecific binding was prevented by saturation with 2% bovine serum albumin (BSA, Sigma Aldrich). Deposited cleaved C4 form was detected with a rabbit anti-human C4 polyclonal antibody (Siemens), an anti-rabbit-HRP antibody conjugate (Sigma Aldrich), addition of TMB (Sigma Aldrich) and a Clariostar plate reader (BMG Labtech). Polyclonal IgM isolated from human serum (Sigma Aldrich) was used as control. Reported values were obtained by normalizing each data set (polyclonal IgM/NHS defined as 100) and by averaging data obtained in replicated independent assays (between 2 to 4); reported errors were the standard deviation of the replicates.

### 2.10 BioLayer Interferometry

All BLI experiments were performed on an OctetRED96e from Pall/FortéBio and were recorded with the manufacturer software (Data Acquisition v11.1). All protein samples were buffer exchanged against 0.01 M Na_2_HPO_4_, 0.0018 M KH_2_PO_4_, 0.137 M NaCl, 0.0027 M KCl at pH 7.4 (Phosphate Buffer Saline, PBS) or 0.025 M Tris, 0.15 M NaCl, 0.003 M KCL at pH 7.4 (Tris Buffer Saline, TBS) with Zeba Spin Desalting columns (Thermo Fisher Scientific) prior to any loading. Commercial AR2G (amine coupling), SA (streptatividin), APS (aminopropylsilane), Protein A, Protein L biosensors (Pall/FortéBio) or lab-made IgM-specific biosensors were tested. The latter ones, mouse or goat anti-μ chain antibodies (Invitrogen) or CaptureSelect anti-IgM nanobody (Thermo Fisher Scientific) were biotinylated using NHS-PEG4-biotin EZ-link kit (Thermo Fisher Scientific) and captured onto SA biosensors. For capture on AR2G biosensors, IgM samples were diluted in 0.01 M Sodium Acetate at pH 4, 4.5, 5, 5.5 or 6 (best capture at pH 6) and C1q in either 0.01 M Sodium Acetate at pH 4, 4.5, 5 or 5.5, 0.01 M MES at pH 6 or 6.5 or 0.01 M HEPES at pH 7 or 7.5 (best capture at pH 7.5); for capture on SA biosensor, IgM or C1q samples were biotinylated using NHS-PEG4-biotin EZ-link kit (Thermo Fisher Scientific) and manufacturer conditions; no chemical treatments were applied before capture on APS, Protein A, Protein L or IgM-specific biosensors. All ligand samples were applied at concentrations between 10 and 50 μg/ml. For association and dissociation, all analyte samples were diluted at concentrations between 10 and 100 nM in TBS complemented with 0.002 M CaCl_2_ and 0.02% Tween-20 as analysis buffer. 0.2 ml of each sample or buffer were disposed in wells of black 96-well plates (Nunc F96 MicroWell, Thermo Fisher Scientific), maintained at 25°C and agitated at 1000 rpm. Biosensors were pre-wetted in 0.2 ml PBS, 0.02% Tween-20 or analysis buffer for 10 min, followed by equilibration in pre-wetting buffer for 120 s. Biosensors were then loaded with each ligand prepared according to the capture chemistry for between 300 s and 600 s, followed by an additional equilibration step of 120 s or more in analysis buffer. In the cases of AR2G biosensors, they were activated by dipping them in a mix of 10 mM N-hydroxysulfosuccinimide (s-NHS) and 20 mM 1-Ethyl-3-3dimethylaminopropyl (EDC) for 300 s prior capture and quenched with 1M Ethanolamine pH8.5 for 300 s after capture. In the case of APS biosensors, they were quenched with 50 μg/ml BSA solution for 600 s. Association phases were monitored during dipping the functionalized biosensors in analyte solutions for between 180 s and 300 s, and the dissociation phases monitored in analysis buffer for between 180 s and 300 s. To assess and monitor unspecific binding of analytes, analyses were performed with biosensors treated with the same protocols but replacing ligand solutions by analysis buffer. Kinetics analyses were performed using Protein L biosensors which were functionalized with each IgM sample diluted at 30 μg/ml for 600 s until reaching a spectrum shift between 5.5 and 7.0 nm. Association phase of plasma C1q was monitored for 300 s with concentrations between 0 and 100 nM and dissociation phase for 600 s. All measurements were performed in replicates with independent IgM loadings. Kinetics data were processed with the manufacturer software (Data analysis HT v11.1). Signals from reference biosensor and zero-concentration sample were subtracted from the signals obtained for each functionalized biosensor and each analyte concentration. Resulting specific kinetics signals were then fitted using a global fit method and 2:1 heterogeneous ligand model. Reported kinetics parameter values were obtained by averaging the values obtained with replicated assays (between 2 to 4) and reported errors as the standard deviation.

## 3 Results

### 3.1 Expression by HEK293F and purification of recombinant IgM in different oligomeric forms

Previously, the production of recombinant IgM models using different mammalian expression systems was intensively studied in stable and transient expression (Wolbank et al., 2003; Vorauer-Uhl et al., 2010; Chromikova et al., 2015b, 2015a; Hennicke et al., 2017, 2019, 2020). Here, we present an additional production system for the two IgM constructs, IgM617 and IgM012, using stabilized HEK293F cell lines. cDNA of H, L and J chains from both models have been individually subcloned in pcDNA3.1(+). To obtain IgM samples in pentameric forms (IgM617-HLJ and IgM012-HLJ), HEK293F cells were transfected with the three H, L and J pcDNA3.1(+) constructs, while to obtain IgM samples in hexameric forms (IgM617-HL and IgM012-HL), cells were transfected with only H and L vector constructs. After selection and culture expansion, recombinant IgMs were purified from culture media using the optimized protocol established by Hennicke *et al*. (Hennicke et al., 2017). As previously observed in other cell lines, the retrieved yields and sample qualities after purification differ from one recombinant IgM to the other one with higher product titers for IgM617 than IgM012. Indeed, the final purified yield from stable HEK293F culture media of IgM617-HLJ or IgM617HL (5 to 10 mg/l of culture supernatant) is 25 to 100 times higher than that of IgM012-HLJ or IgM012-HL (less than 0.1 mg/l of culture supernatant). Furthermore, the production yield and sample quality of IgM617 appear reproducible from different cell cultivations and are far more variable for IgM012s and often, samples suffer from low amounts of product and homogeneity.

### 3.2 Biophysical characterization of the oligomeric forms of recombinant IgMs

The oligomeric distribution and quality of purified IgM samples expressed by HEK293F cell lines were investigated with biochemical, biophysical and structural methods when feasible.

#### 3.2.1 SDS-PAGE

Semi-native PolyAcrylamide Gel Electrophoresis (PAGE) adapted from Vorauer *et al*. (Vorauer-Uhl et al., 2010) shows that purified IgM617-HLJ migrates as a single and homogeneous band while IgM617-HL migrates in a more heterogenous manner with two major bands in the highest molecular weight range, suggesting thus the presence of different high oligomeric states in the absence of J chain (**Figure 1A**). Interestingly, and while IgM012-HLJ and IgM012-HL behave similarly to IgM617 constructs, noticeable additional bands at molecular weights below 150 kDa are still observed, suggesting the presence of lower oligomeric states (**Figure 1B**) similarly to what has been described by Chromikova *et al*. (Chromikova et al., 2015b) and Hennicke *et al*. (Hennicke et al., 2020) for the production of IgM617 and IgM012 pentamers in CHO DG44 or HEK239E cell lines.

**Figure 1.**
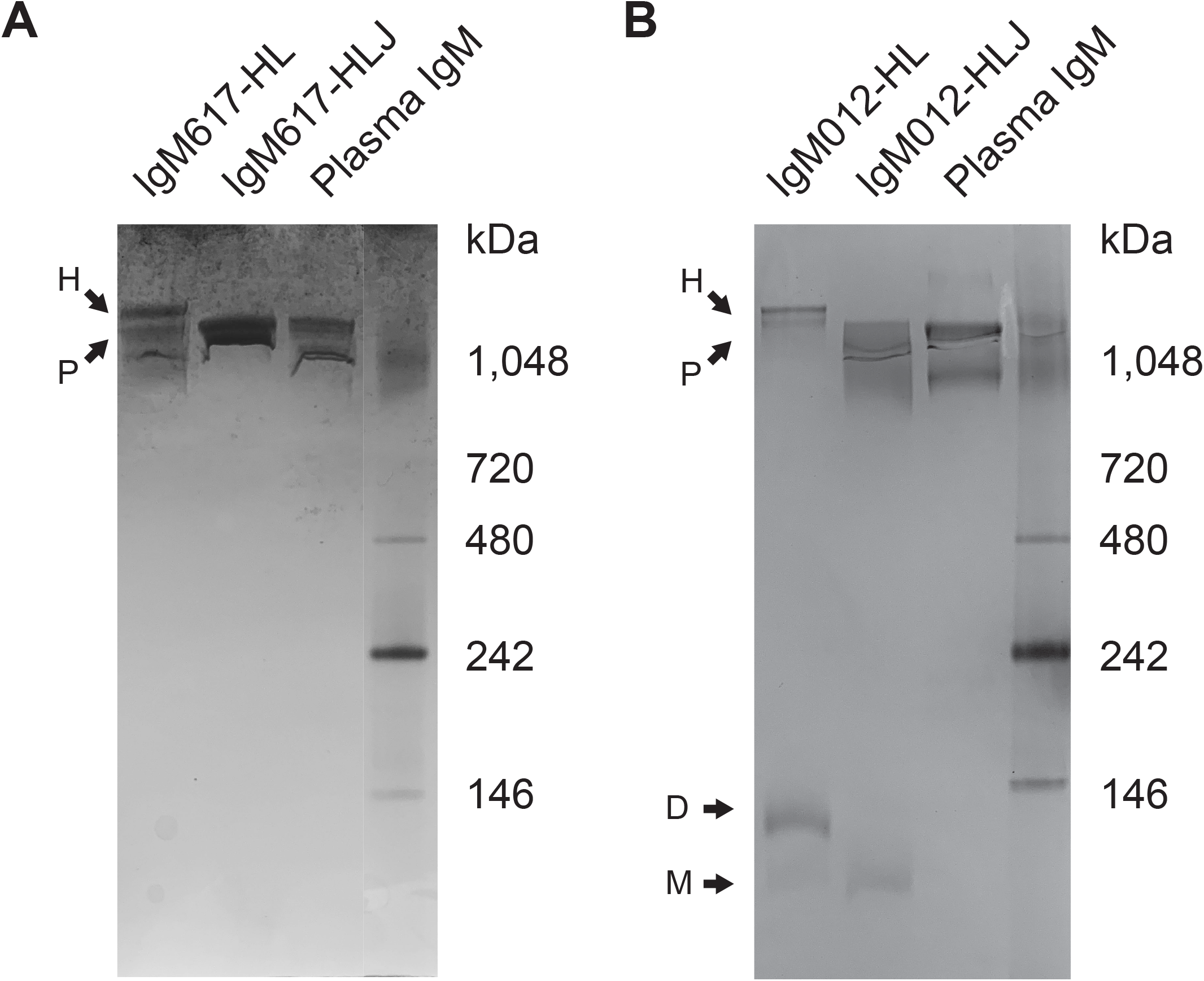
Semi-native PolyAcrylamide Gel Electrophoresis (PAGE) analysis of purified IgMs samples from HEK293F expression. Native PAGE gels followed by Coomassie Blue staining were run to identify **(A)** IgM617-HLJ and IgM617-HL and **(B)** IgM012-HLJ and IgM012-HL polymer distributions after SEC purification. IgM purified from plasma (Antibodies-onlines) are taken as makers along with Native Markers. Suspected hexamers (H), pentamers (P), dimers (D) and monomers (M) are indicated with arrows.

#### 3.2.2 SEC-MALLS and AUC

To further characterize IgM617-HLJ and IgM617-HL homogeneities, Multi-Angle-Light Scattering coupled to Size Exclusion Chromatography (SEC-MALLS) and sedimentation velocity Analytical Ultra-Centrifugation (sv-AUC) were performed.

SEC-MALLS analyses gave a homogeneous sharp peak of about 895 kDa for IgM617-HLJ which falls in the range of theoretical molecular weight of the glycosylated pentamers (891 kDa of amino acids + 10 to 15% of glycosylation) (**Table 1** and **Figure 2B**). Surprisingly, a single mass of 947 kDa is also observed for IgM617-HL (**Table 1** and **Figure 2A**), while the construct was anticipated by SDS-PAGE analysis to be a mixture of pentamers and hexamers (see above). The observed mass may likely correspond to average molecular weights from the different glycosylated oligomers (hexamers: 1050 kDa; pentamers without J chain: 875 kDa + 10 to 15%). They might thus co-elute as a single peak from SEC and the resolution of this method is not sufficient to separate hexamers and pentamers. To further explore sample heterogeneities, sv-AUC experiments were performed. Migration in the velocity field of IgM617-HLJ samples showed one main homogeneous peak at a sedimentation coefficient of about 19 S (more than 75 %) and minor peaks at 12 S (12 %) and 6 S (9 %), revealing the presence of a majority of pentamers but also presence of lower molecular weight oligomers (**Table 1** and **Figure 2D**). Migration of IgM617-HL samples appeared to be even more heterogeneous with two main peaks at about 21 S (between 60 to 70 % depending of lots) and about 17.5 S (25 to 30 %), which may correspond to hexamers and pentamers, respectively; and two minor peaks at 15.0 S (5 %) and below 5 S (3 to 10 %), also lower molecular weight oligomers (**Table 1** and **Figure 2C**). Our AUC data on recombinant IgMs are perfectly in agreement with the sedimentation coefficients originally measured by Eskeland *et al*. in 1975 (Eskeland and Christensen, 1975) on purified IgM from patient sera with or without J chain.

**Table 1.** Summary of theoretical and experiment molecular weights of sedimentation coefficients determined by AUC. The theorical peptide molecular weights are calculated based on the amino-acid primary sequences of the subcloned IgM chains and using Protparam (Gasteiger et al., 2005). Experimental molecular weights were determined by SEC-MALLS and Mass Photometry, coefficient sedimentation by AUC.

| IgM | Oligomers | Theoretical (kDa) | SEC-MALLS (kDa) | AUC (S) | Mass Photometry (kDa) |
| --- | --- | --- | --- | --- | --- |
| IgM617-HL | Hexamers | 1050 | 947 | 21.0 S | 1250 ± 54 |
|  | Pentamers | 875 |  | 17.5 S | 1050 ± 43 |
|  | Tetramers | 700 | - | 15.0 S | 841 ± 37 |
|  | Dimers | 175 | - | 5.0 S | 162 ± 12 |
| IgM617-HLJ | Pentamers | 891 | 895 | 19 S | 998 ± 6 |
|  | Tetramers | 700 | - | 12 S | - |
|  | Dimers | 175 | - | 6 S | - |
| IgM012-HL | Hexamers | 1034 | n.d. | n.d. | 1166 ± 21 |
|  | Pentamers | 862 | n.d. | n.d. | 1052 ± 34 |
|  | Tetramers | 690 | n.d. | n.d. | - |
|  | Dimers | 172 | n.d. | n.d. | 150 ± 21 |
| IgM012-HLJ | Pentamers | 878 | n.d. | n.d. | 944 ± 32 |
|  | Tetramers | 690 | n.d. | n.d. | - |
|  | Dimers | 172 | n.d. | n.d. | 162 ± 14 |

(n.d: not determined)

**Figure 2.**
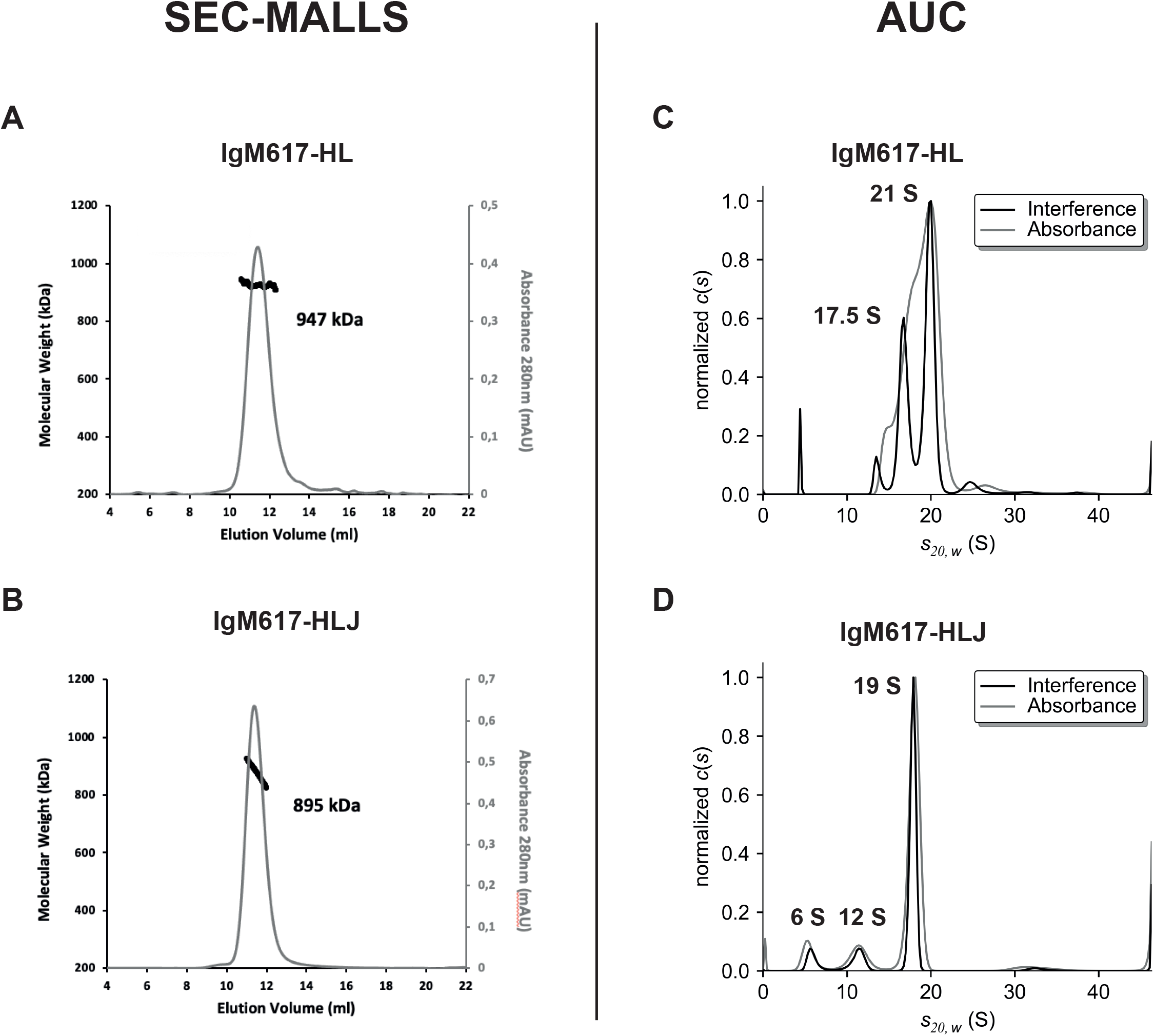
Size Exclusion coupled to Multi-Angle Light Scattering (SEC-MALLS) and Analytical UltraCentrifugation (AUC) analysis of purified IgM617 samples from HEK293F expression. On the left panels, the chromatograms show the elution profile of the purified **(A)** IgM617-HL and **(B)** IgM617-HLJ monitored by excess refractive index (left ordinate axis) and the molecular weight as bold line (right ordinate axis) derived from MALLS, refractometry and UV-Vis measurements. The estimated average molecular weights are indicated on the graphs. On the right panels, the sedimentation distributions of **(C)** purified IgM617-HL and **(D)** IgM617-HLJ. Calculated and corrected sedimentation coefficients S_*20,W*_ are obtained as in Materials and Methods and are indicated on the graphs.

#### 3.2.3 Mass Photometry

To assess more precisely their oligomeric states and experimental molecular masses, IgM samples were further analyzed using an emerging technique enabling accurate native mass measurements of single molecules in solution, the Mass Photometry (MP) (Sonn-Segev et al., 2020). A single population with an average mass of 998 +/-6 kDa (mean +/-SD over replicates) can be observed for IgM617-HLJ (**Table 1** and **Figure 3B**). The experimental mass matches the mass of fully glycosylated pentamers (891 kDa + 10 to 15% glycosylation). By contrast, IgM617-HL data show several mass populations: 3 main populations at 1250 +/-54 kDa, 1050 +/-43 kDa, and 841 +/-37 kDa (**Table 1** and **Figure 3A**) that may correspond to hexamers (1050 kDa + 10 to 15% glycosylation), pentamers (875 kDa + 10 to 15%) and tetramers (700 kDa + 10 to 15%), respectively. Minor populations can also be observed and might correspond to lower oligomeric protomer states. This is in agreement with our sv-AUC observation of a mixture of IgM617-HL oligomers (see above). However, the oligomeric distributions appear different and might be accountable for by the usage of different lots of IgM617-HL for the measurements but also by the methods used. IgM012-HLJ appears with a quite homogeneous experimental mass of 944 +/-32 kDa, corresponding to pentamers (878 kDa + 10 to 15%) although a broad mass distribution is still observed and an additional population at low molecular weight is present and may correspond to the lower oligomeric states (**Table 1** and **Figure 3D**). Finally, IgM012-HL behaves similarly to IgM617-HL with 2 main populations at 1166 +/-21 kDa and 1052 +/-34 kDa corresponding to hexamers (1035 kDa + 10 to 15%), and pentamers (862 kDa + 10 to 15%), but low oligomeric states can also be observed (**Table 1** and **Figures 3C**).

**Figure 3.**
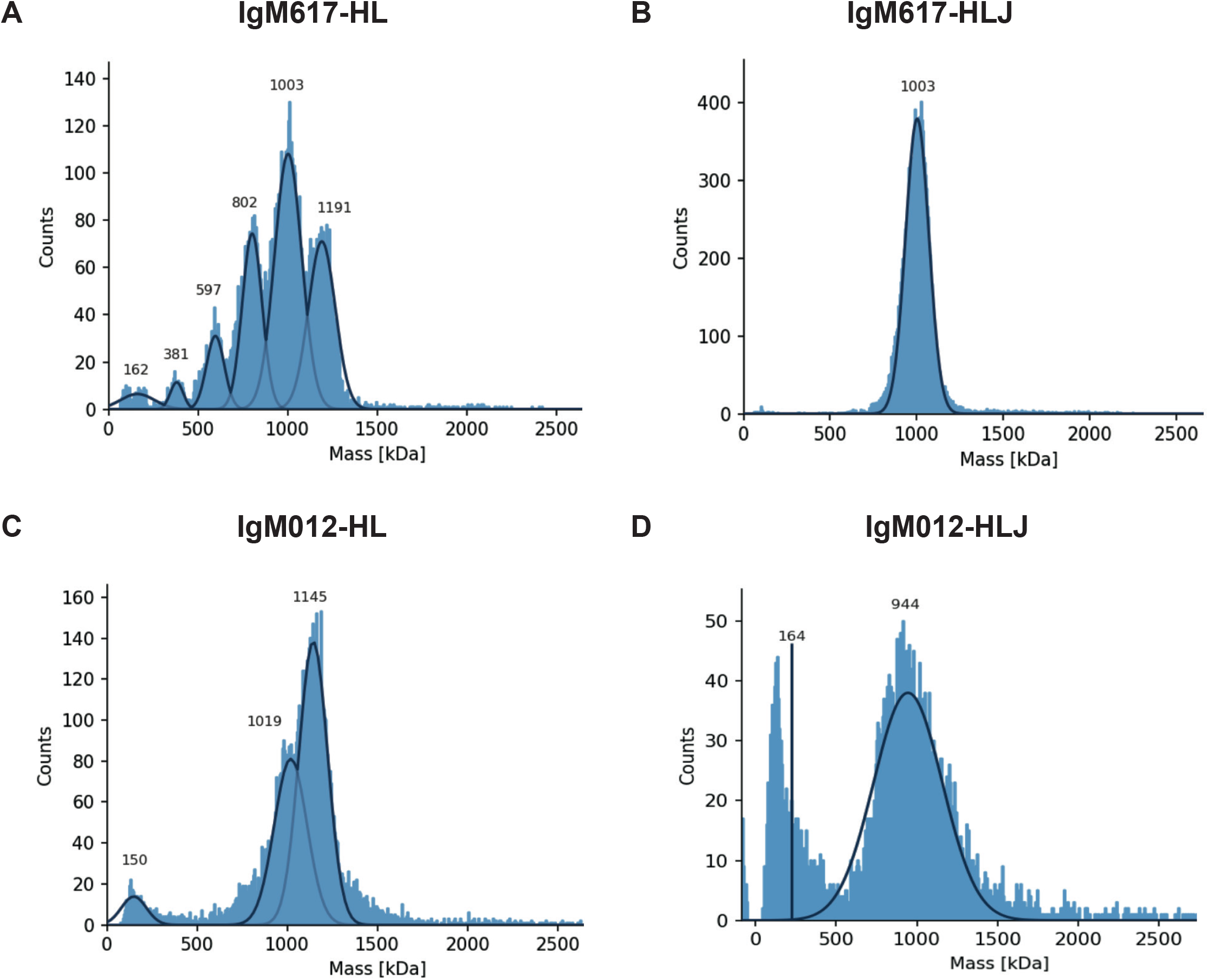
Mass Photometry analysis of recombinant IgMs produced in HEK293F. Histograms show the population distributions of purified IgMs. Shown analyses are representative of replicate experiments: **(A)** IgM617-HL, **(B)** IgM617-HLJ, **(C)** IgM012-HL and **(D)** IgM012-HLJ. Estimated molecular weights for the shown experiments are indicated on each graph.

#### 3.2.4 Transmission Electron Microscopy

The structural integrity and oligomeric states of IgMs produced by HEK293F were also examined and confirmed with negative staining Transmitted Electron Microscopy (TEM). IgM617-HLJ (**Figure 4B**) and IgM012-HLJ (**Figure 4D**) produced in HEK293F exhibit very similar structural characteristics as previously observed for pentamers produced in HEK293E and CHO DG44, with a central circular core with projecting flexible Fab units in a star-shaped manner as well-known for IgMs isolated from human serum. As expected from biophysical data described above, IgM617-HL (**Figure 4A**) and IgM012-HL (**Figure 4C**) present distinct types of particles: some with a symmetric shape and six arms, confirming the hexameric structural features of HL samples in addition to the pentamer ones, which is also confirmed by particles with an asymetric shape to which five arms can be attributed. As a remark, IgM molecules with lower oligomeric states observed with Mass Photometry and AUC could not be easily identified in TEM micrographs.

**Figure 4.**
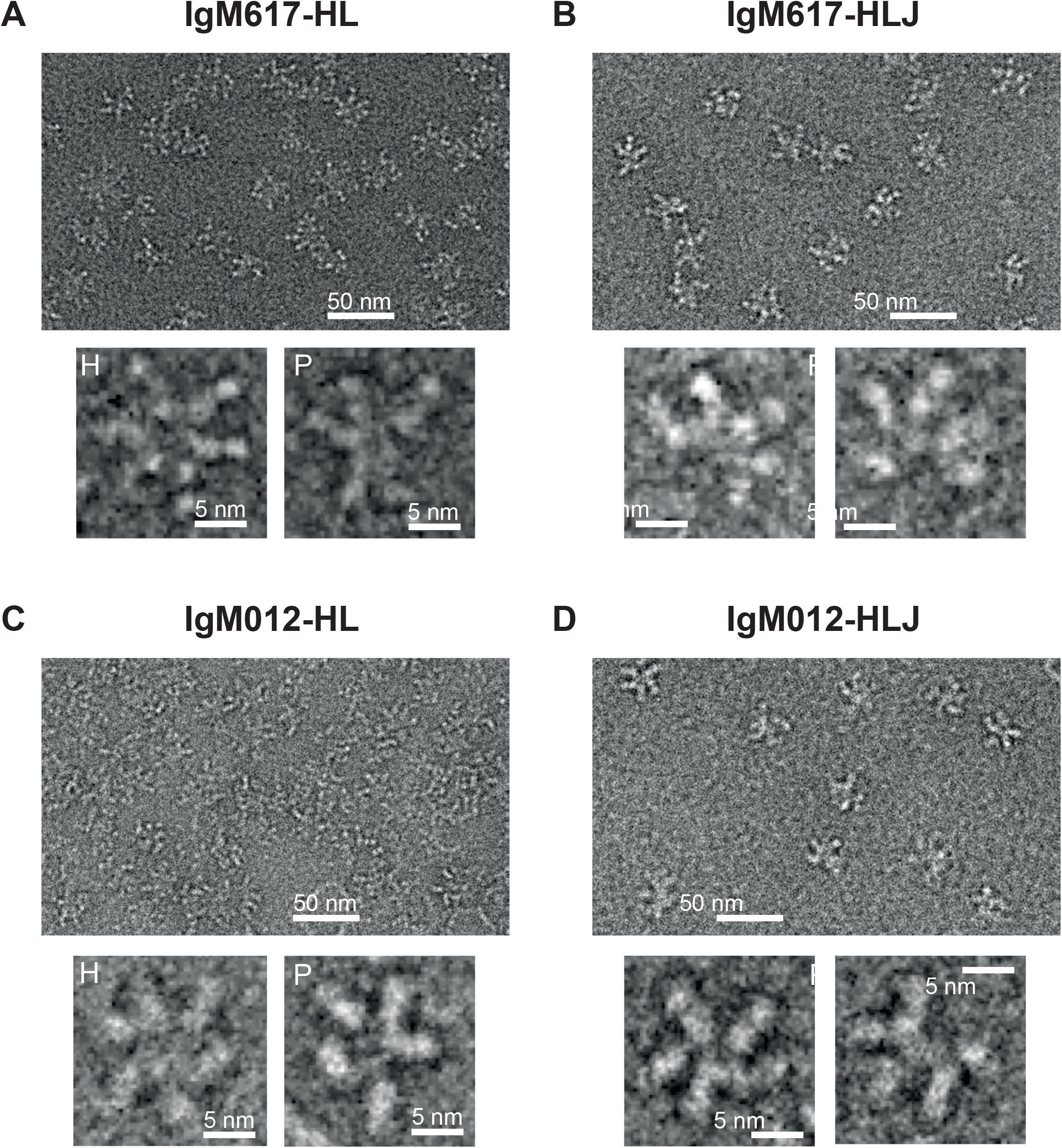
Non-processed images of negative stain transmission electron microscopy images of recombinant IgMs produced in HEK293F. Representative fields of particles with a 50 nm scale bar on top of each panel and magnified views of some individual molecules are shown on the lower part of each panel: **(A)** IgM617-HL, **(B)** IgM617-HLJ, **(C)** IgM012-HL, **(D)** IgM012-HLJ. Hexamers are denoted with H and Pentamers with P.

Taken together, the usage of several biophysical characterization methods allows to show that the retrieved production yields and protein quality in terms of homogeneity and oligomer distribution differ significantly depending on individual recombinant IgM construct. Our presented data confirmed expression of a majority of pentameric IgM in stable HEK293F cell lines transfected with the 3 IgM genes. When only H and L chain genes are transfected to obtain stable HEK293F cells, we also observed the formation of hexamers in addition to pentamers, likely missing the J chain. However, hexamer ratio appears to be variable and appearance of incomplete polymers can be observed in the HL sample preparations.

### 3.3 C1q-dependent complement activation by recombinant oligomeric IgM forms produced by HEK293F cells

The capacity of the different recombinant IgM preparations to activate CP was analyzed using our in-house *in vitro* Enzyme-Linked Immunosorbent Assay (ELISA) based on the detection of C4b fragment deposition after serum cleavage of C4 by the C1 complex bound to coated IgM molecules (Bally et al., 2019; Hennicke et al., 2020). Assays with C1q-depleted normal human serum and C1q-reconstituted serum were used as controls for the C1q/IgM interaction dependency. Polyclonal IgMs purified from human plasma (pIgMs) were used as positive control and standard. As observed with our previous IgM617 and IgM012 productions (Hennicke et al., 2020), no meaningful differences are observable in the C4b deposition yields between the different coated IgM samples, either serum-derived or recombinant. Thus, the polymer distributions, might not influence the ability of IgMs to activate the proteolytic complement cascade through C1 when coated to the ELISA surfaces (**Figure 5**).

**Figure 5.**
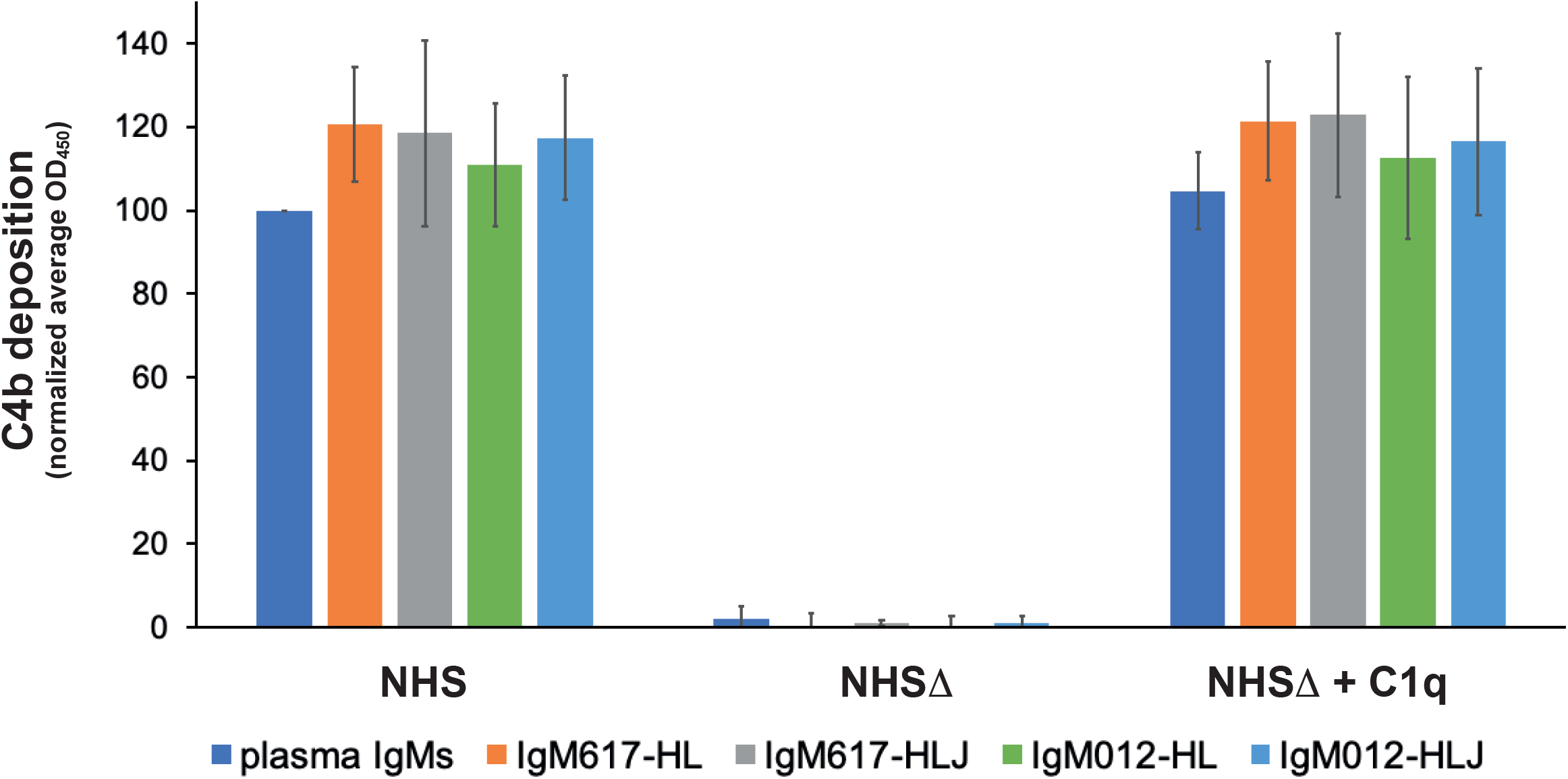
Complement activation by the different purified IgMs constructs expressed in HEK293F. Recombinant IgMs and IgMs purified from plasma were coated on microplate wells and incubated with either Normal Human Serum (NHS), C1q-depleted NHS (NHSΔ) or reconstituted NHS (NHSΔ + C1q). C1 activity was monitored via C4b deposition. Reported values are average of replicated (2 to 4) and normalized experiments (100% = IgM plasma/NHS). Errors are obtained as standard deviation between independent replicates.

### 3.4 C1q binding kinetics to IgMs measured by BLI

In order to characterize the binding kinetics of IgMs to C1q, we developed new protocols using a label-free optical method based on the Reflectometric Inference Spectroscopy (RIfS) (Hänel and Gauglitz, 2002; Gauglitz, 2020) and its setup known as BioLayer Interferometry (BLI) (Abdiche et al., 2008; Sultana and Lee, 2015) on an OctetRED96e instrument. During the course of this study, plasma purified C1q and plasma-derived IgM preparations or recombinant IgM constructs produced in HEK293F, complemented with IgMs produced in CHO-DG44 and HEK293E (Hennicke et al., 2020) were used. Real-time detection of binding events at the surface of biosensors enables two strategies: (i) using C1q captured as ligand to the biosensor surfaces and the IgMs as analytes or (ii) IgMs captured as ligands to the biosensor surfaces and C1q as the analyte.

The first strategy has been evaluated with C1q as ligand either amine coupled onto AR2G biosensors or captured by Streptavidin (SA) biosensors after biotinylation. However, although high C1q densities can be reached (3 to 7 nm spectral shift), no binding responses can be retrieved with plasma IgM samples (**Supplementary Figure S1**).

In the second strategy, we tested several different commercial biosensors (AR2G, SA, APS, Protein A and Protein L) and lab-made IgM-specific biosensors (mouse or goat anti-μ chain antibodies or CaptureSelect anti-IgM nanobody coated on SA biosensors). All biosensors could be functionalized with plasma-derived polyclonal IgMs at different high levels depending of the capture chemistry (between 1 to 7 nm spectral shift) (**Supplementary Figure S2 left panels, Figure S3B**). Since Protein A is specific to Fcγ regions of IgG but also to VH_3_ subfamilies of Ig, both monoclonal recombinant IgM617 and IgM012 have been tested but no capture signal could be retrieved for this type of biosensors although IgM617 belongs to the VH_3_ subfamily. Unspecific binding of C1q was also evaluated for all biosensor types: without any captured IgM, most of them showed either low or no unspecific signals from C1q, contrary to what was previously reported (Zhou et al., 2018) (**Supplementary Figures S2 right panels**). Finally, either no or too weak C1q binding signals can be retrieved from most of the IgM-functionnalized biosensors, making them not applicable kinetics and affinity determination (AR2G, SA, Protein A, APS, anti-μ chain antibodies) (**Supplementary Figure S2 right panels**). Only protein L biosensors show both limited signals of unspecific C1q binding (**Supplementary Figure S3A**) and measurable kinetics signals on the IgM-functionalized biosensors.

Thus, pIgMs, IgM617 and IgM012 constructs were stably absorbed on Protein L biosensors until reaching saturation (4 to 7 nm shift) (**Supplementary Figure S3B**) and kinetics analyses were then performed by dipping the IgM-functionalized biosensors in C1q concentration series ranging from 3.13 to 100 nM with 2-fold dilution (**Figure 6**).

**Figure 6.**
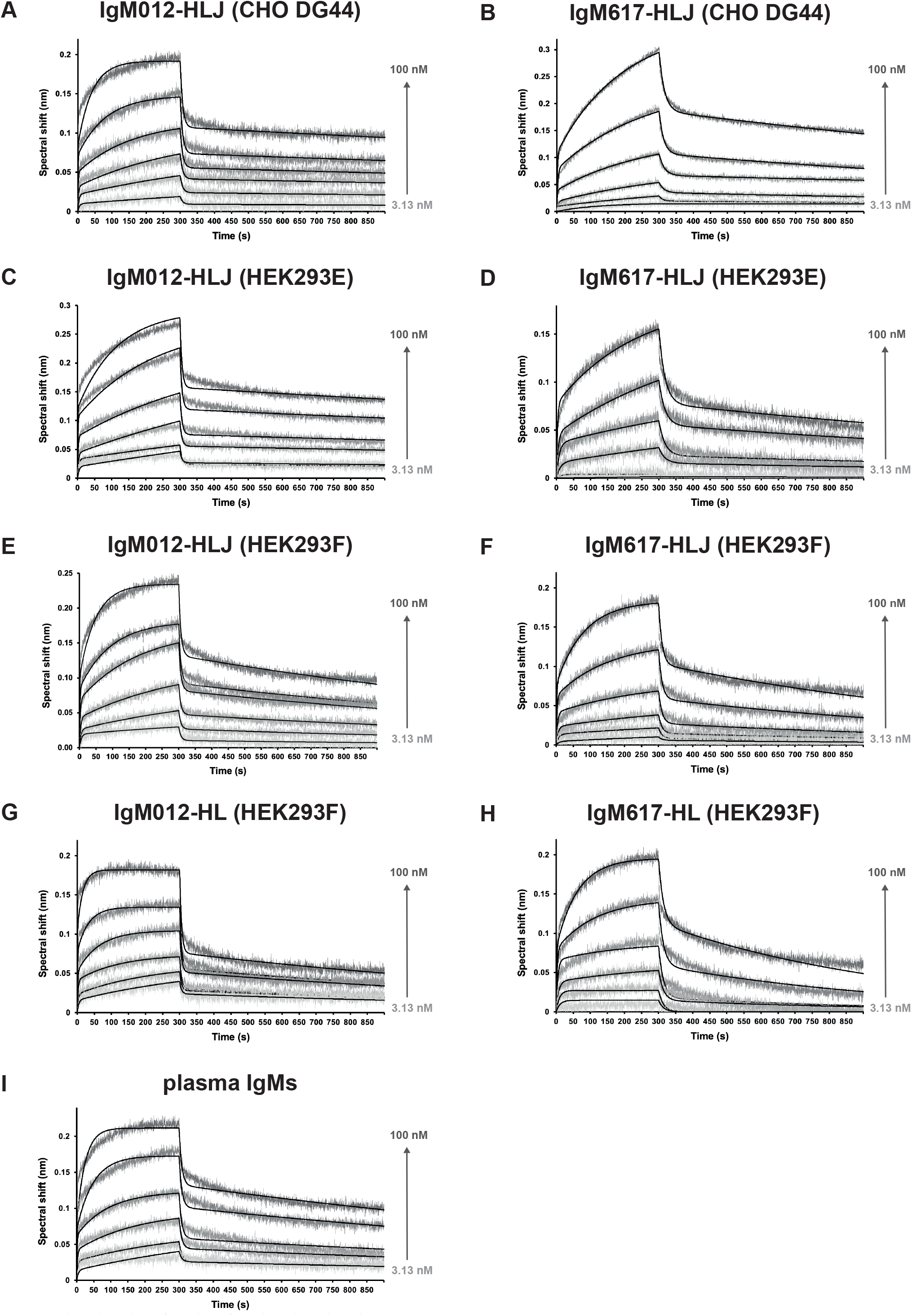
Kinetics analysis of the interaction between C1q from plasma and the different purified and immobilized full IgMs expressed in mammalian expression hosts. IgMs were captured on Protein L and functionalized biosensors were dipped in wells containing C1q at different concentrations (3.13, 6.25, 12.5, 25, 50, 100 nM). The binding signals (grey-scaled sensorgrams) were obtained by subtracting the signals from empty protein L biosensor and from zero-concentration samples. Fitted curved are depicted as black lines and were obtained by global fitting using a 2:1 heterogenous ligand model. Shown kinetics analysis are representative of each binding experiment: **(A)** IgM012-HLJ expressed by CHO DG44, **(B)** IgM617-HLJ by CHO DG44, **(C)** IgM012-HLJ by HEK293E, **(D)** IgM617-HLJ by HEK293E, **(E)** IgM012-HLJ by HEK293F, **(F)** IgM617-HLJ by HEK293F, **(G)** IgM012-HL by HEK293F, **(H)** IgM617-HL by HEK293F, **(I)** IgMs purified from plasma.

C1q binds to all IgMs in a biphasic manner, with two distinguishable fast and slow binding rates, and the 1:1-Langmuir model could not be applied to determine interaction kinetics parameters. Thus, the only fitting model available to take into account both kinetics components was the 2:1-heterogeneous-ligand model (**Figure 6**). Polyclonal IgMs extracted from plasma and all recombinant monoclonal IgMs show affinities in the nM scale, ranging from about 1 nM to 8 nM for K_D1_ and from 4 to 26 nM for K_D2_, depending on the sample (**Table 1**). Although the differences may not be significant, IgM012s seems to possess slightly better affinity for C1q (K_D1_ = 1 to 3 nM, K_D2_ = 4 to 10 nM) than IgM617s (K_D1_ = 6 to 8 nM, K_D2_ = 7 to 26 nM). They thus behave more similarly to pIgMs (K_D1_ = 2.3 nM, K_D2_ = 6.5 nM) than IgM617s. It has to be noted that for all IgM samples, both distinguishable slow and fast kinetics lead to affinities in the same nM order of magnitude. No significant differences in kinetics, which could be related to the host cell line, can be observed either for IgM617 or IgM012 models. Finally, although small differences can be noted in kinetics rate values when comparing HL and HLJ constructs, it is difficult to attribute them solely to the presence of hexameric forms, since such variations are also observed either between HLJ constructs or between different preparations of the very same recombinant IgM construct (**Table 1**).

## 4 Discussion

In the presented study, we recombinantly produced and purified two IgMs models, IgM617 and IgM012 (Hennicke et al., 2017, 2019, 2020) in different oligomeric states and fully characterized their biochemical and functional quality attributes using combined biophysics techniques : (i) AUC, SEC-MALLS, MP and TEM to determine their oligomeric states and polymer distributions, (ii) *in vitro* complement activation based on ELISA to evaluate their functional capacities. Moreover, we developed a new protocol using a label-free and real-time detection technique, the BLI, to quantify the kinetics of the interaction between IgMs and the recognition and activating protein of the classical complement pathway, C1q.

One challenge for studying IgM and developing potential therapeutic products is the manufacturing and characterization of recombinant molecules of high quality in terms of Ig productivity, homogeneous oligomerization status, purity and biological biomimetic function. In the past years, several methods have been tested and in particular both IgM617 and IgM012 have been taken as IgM models to develop proof-of-concepts of recombinant production (Vorauer-Uhl et al., 2010; Mader et al., 2013; Chromikova et al., 2015b; Hennicke et al., 2017, 2019, 2020). In the present study, we complemented biophysical sample analysis method panel by using SEC-MALLS (**Figure 2**), sv-AUC (**Figure 2**) and the new MP (**Figure 3**) to determine the oligomeric distribution of recombinant IgM samples produced in HEK293F cell line. We can thus confirm that IgM617-HLJ model can be produced as very homogeneous full pentamers when H, L and J chain genes are transfected in HEK293F for stable expression, while IgM012-HLJ retains heterogeneities with significant amounts of lower oligomers. Despite these challenging behaviors, both IgM pentamers produced in HEK293F retain their structural integrity of five-branches-star-shaped particles (**Figure 4**) as shown with negative staining TEM images and functional qualities as demonstrated with complement activation assays (**Figure 5**). One explanation for the challenging behavior of IgM012 in recombinant expression compared to IgM617 and in general to other IgM constructs studied so far, may come from its origin: it is a class switch of the human monoclonal anti-HIV-1 antibody 2G12 (Wolbank et al., 2003). Indeed, the 2G12 Fab crystal structure revealed an unusual VH-VL domain arrangement for Fabs where two of its Fabs assemble as a dimer with interlocked VH domain swapping, instead of the usual VH-VL monomeric assembly (Calarese et al., 2003). The VH domain exchange has been demonstrated to be mandatory for this IgG to target and to recognize the specific high-mannose cluster of the glycan shield of HIV-1 (Doores et al., 2010). One can easily hypothesize that IgM012 might possess such similar domain swapping of its Fabs and this might constrain IgM assembly more than the regular one in which Fabs behave independently. These structural constraints might not be favorable and might impair a proper expression of the IgM012 recombinant constructs by the different cell lines used so far.

With this study, we show that, by stably transfecting HEK293F cells with only the H and L constructs, IgM617-HL, and to a lesser extend IgM012-HL, hexamer expression is made possible using HEK293F. It was well established that the J chain expression influences the formation of the pentameric and hexameric IgM forms either *in vivo* or *in vitro*. Randall *et al*. in 1992 first demonstrated the pentamer/hexamer ratio secretion regulation in B cells by J chain gene expression with the secretion of a majority of hexamers and a minority of pentamers by cells lacking the J chain gene (Randall et al., 1992). However, recombinant expression of only hexamers by J-chain deficient cells remains challenging. Although hexamers were found to be expressed by recombinant systems, a majority of pentamers is found to be secreted by mouse hybridoma cell lines (Wiersma et al., 1998; Collins et al., 2002) or CHO-K1(Gilmour et al., 2008). Interestingly, only Azuma *et al*. succeeded to obtain a majority of hexamers by using CHO-DG44 that expressed 20 times more hexamers that the CHO co-transfected with the J chain gene (Azuma et al., 2007). Here, we confirm that recombinantly producing homogeneous hexameric forms of IgM remains difficult. Although IgM617 and IgM012 models can be expressed and secreted as hexamers by stable HEK293F cell lines as shown by our different biochemical, biophysical and structural analyses, our data also show that both sample types are not homogeneous: they present a large population of pentamers likely lacking the J chain, but also traces of lower molecular weighted oligomers, likely tetramers and dimers, in the purified IgM samples. Moreover, the ratio between the distinct IgM oligomeric states appears not only variable from an IgM model to the other one but also between different production runs of the same IgM model (data not shown). Additionally, our data from our surface-based C1q-dependent complement activation assays do not demonstrate higher activities of IgM samples enriched with hexamers (**Figure 5**). This contrasts with previous hemolytic assays, which demonstrated the higher efficacy of hexameric IgM to induce complement-dependent cytolysis compared to pentameric IgMs (Randall et al., 1990; Collins et al., 2002). As pointed previously, the ELISA-like setup and the end-point measurement of C4 deposition might not be sensitive and resolutive enough to characterize differences in IgM abilities to activate the first activation step of the complement (Hennicke et al., 2020).

For the first time, we introduced a new method to characterize the binding kinetics and affinities of C1q to IgM antibody isotypes using BLI. Optical and surface-based methods such as SPR and BLI are popular methods to characterize the binding properties of all antibody classes to antigens, but much less to Ig effectors such as the complement molecule, C1q. Patel *et al*. in 2015 (Patel et al., 2015) and Jovic *et al*. in 2019 (Jovic and Cymer, 2019) have successfully characterized the binding responses of C1q to the different IgG subtypes using traditional SPR methods and Zhou *et al*. in 2018 (Zhou et al., 2018) using the recent BLI technology. Concerning the IgM/C1q interaction properties, a single study has been published by Bally *et al*. in 2019 (Bally et al., 2019) who measured, using SPR, the binding kinetics between IgM fractions purified from human plasma and native C1q, as well as recombinant C1q and a few critical mutants.

Several experimental setups have been tested in our study to define the conditions allowing reliable binding measurements between either IgM purified from human plasma or different recombinant IgM constructs and native C1q purified from human plasma. Surprisingly and in contrast to the SPR data obtained so far when C1q is captured at the biosensor surface (Bally et al., 2019), no interaction with IgMs was detectable with the BLI method (**Supplementary Figure S1**). However, we could observe binding of the catalytic tetramer C1r_2_Cs_2_ to immobilized C1q with expected affinities in the nanomolar range (**Supplementary Figure S4**). These results indicate that immobilized C1q can retain certain binding activities towards its partners but that the immobilization strategy to the BLI biosensor either directly or via selective biotinylation of primary amines of C1q can greatly affect its binding capacities.

In the course of optimizing the BLI protocols, capturing IgMs on Protein L biosensor and using C1q as analyte has proven to be the most efficient strategy since only low or acceptable unspecific C1q binding can be detected at the used concentration range (**Supplementary Figure S3**) and specific concentration dependent binding signals can be measured (**Figure 6**). Our method meets the ones already employed to characterize the complexes between C1q and recombinant IgGs, for which Protein L coated sensors have been used to capture immunoglobulins for SPR or BLI, although Streptavidin and biotin fusion have also been proven to be suitable (Zhou et al., 2018). Unfortunately, protein L was the only successful immobilization method for functionalizing BLI biosensors with IgMs. Indeed, the binding of protein L to Igs is specifically restricted to certain kappa light chains, which limits the application of the method described here to IgMs containing this subtype of L chain, such as our models. Surprisingly, binding of C1q can be measured in those conditions and without any specific antigen bound to IgMs. Similarly, complement activation can be quantified without antigen with solid phase methods such as ELISA like techniques as shown by our presented and previous data (Bally et al., 2019; Hennicke et al., 2020), while antigen is required in solution methods such as erythrocyte or liposome lysis (for assays review in Harboe et al., 2011). Indeed, it is thought that C1q binding to IgM and subsequent activation of complement cascade are made possible only after IgM binding to specific antigen which induces large structural changes in IgM quaternary structure and the exposure of the cryptic binding sites for C1q globular heads (Sharp et al., 2019). One possible explanation would be that immobilizing IgMs over an *in vitro* surface might somehow provoke the necessary structural feature exposures. In particular for BLI experiments for which only IgM captures via Protein L allow C1q binding, the explanation might come from the binding properties of Protein L to Ig. As shown by the 3D structure of the complex between Protein L and Fab, one Protein L domain bridges two Fab domains (Graille et al., 2001). Although this property may not strictly mimic antigen binding, it might be enough to induce an IgM conformation allowing complex formation with C1q.

As expected, the measured affinities of C1q for IgMs with BLI fall within the nM range (**Table 2**). No significant differences in affinities and kinetics rates could be observed between polyclonal IgM purified from plasma and the different monoclonal recombinant IgMs. Together with the complement assays, which also show no differences between IgM samples, our BLI data emphasize that recombinant IgM preparations retain similar abilities as physiological IgMs to bind and to activate the CP *in vitro*. Our data are very consistent with SPR data obtained by Bally *et al*. (Bally et al., 2019). However, the kinetics behavior appears to be different since a 2:1 heterogenous model has been applied to fit our BLI binding data (**Figure 6**), while a 1:1 Langmuir model is sufficient to interpret SPR data. One can argue that the biphasic kinetics behavior might be raised by the heterogenous quality of native or of our recombinant IgM samples. However, no difference is observed between the most homogeneous constructs (*i*.*e*. IgM617-HLJ) and the other ones. Thus, the differences observed between SPR and BLI data likely originate from the used methods and strategies: the used functionalized sensors have different capture chemistries and captured molecules (Amine coupled C1q in case of SPR, Protein L/IgMs in case of BLI) and different molecules are used as analyte (IgM in case of SPR, C1q in case of BLI). The question will remain unanswered since no binding could be detected with SPR, capturing IgMs on Protein L sensors and using C1q as analyte (data not shown). At last, BLI measurements of the binding between IgMs and C1q might reveal intrinsic behaviors of the complex formation between those highly oligomeric and flexible molecules. It is expected that single globular region of C1q may have a lower affinity for Igs than full C1q, as well as IgM protomer for C1q than full IgM. Thus, the apparent and necessary affinities for the C1q/IgM complex formation rely on the specific multivalency that enhance the binding with an avidity effect. Furthermore, a certain liability of the C1q/Ig complex formation has been observed: not all gC1q bind the IgG hexamers at the same time (Ugurlar et al., 2018), although the same behavior has not been described for IgMs (Sharp et al., 2019). Finally, because of the very high flexibility of IgM and C1q molecules, structural arrangements and stabilization are highly expected during the binding events. All taken together, these molecular characteristics would induce very complicated and heterogenous binding kinetics for which any mathematical fitting model would not be sufficiently formulated and easily applied to consider all possible behaviors.

**Table 2.** Kinetics and affinity constants of C1q binding to the different purified recombinant IgMs expressed in mammalian expression systems. Values are obtained after global fitting of the binding signals (see Figure 5) and averaging from replicates (2 to 4). Affinity constants (K_D1_ and K_D2_) are obtained from the ratio between kinetics parameters (k_d1_/k_a1_ and k_d2_/k_a2_). Standard errors are obtained as standard deviation between replicates.

| **IgM** | **Source** | **Construct** | $k_{a1}$<br>10<sup>5</sup>/Ms | $k_{d1}$<br>10<sup>-4</sup>/s | **$K_{D1}$**<br>**10<sup>-9</sup> M** | $k_{a2}$<br>10<sup>7</sup>/Ms | $k_{d2}$<br>10<sup>-1</sup>/s | **$K_{D2}$**<br>**10<sup>-9</sup> M** |
| --- | --- | --- | --- | --- | --- | --- | --- | --- |
| **Total IgM** | **plasma** |  | 3.04 ± 0.47 | 7.02 ± 1.63 | **2.31 ± 0.09** | 2.00 ± 0.90 | 1.30 ± 0.60 | **6.52 ± 1.34** |
| **IgM617** | **CHO DG44** | **HLJ** | 0.78 ± 0.17 | 6.20 ± 1.02 | **8.00 ± 0.31** | 0.57 ± 0.44 | 0.74 ± 0.08 | **12.30 ± 3.87** |
|  | **HEK293E** | **HLJ** | 0.92 ± 0.16 | 6.26 ± 1.77 | **6.82 ± 0.11** | 0.86 ± 0.49 | 0.59 ± 0.33 | **6.90 ± 1.13** |
|  | **HEK293F** | **HLJ** | 1.59 ± 0.76 | 11.30 ± 3.04 | **7.10 ± 1.06** | 0.44 ± 0.05 | 1.15 ± 0.53 | **26.4 ± 2.89** |
|  | **HEK293F** | **HL** | 1.66 ± 0.50 | 9.25 ± 4.84 | **5.59 ± 1.02** | 0.96 ± 0.38 | 0.96 ± 0.47 | **9.97 ± 2.00** |
| **IgM012** | **CHO DG44** | **HLJ** | 2.22 ± 0.08 | 2.57 ± 0.37 | **1.16 ± 0.01** | 2.04 ± 0.31 | 1.34 ± 0.13 | **6.54 ± 0.11** |
|  | **HEK293E** | **HLJ** | 1.64 ± 0.35 | 3.57 ± 0.33 | **2.18 ± 0.06** | 1.84 ± 0.40 | 1.57 ± 0.15 | **8.51 ± 0.24** |
|  | **HEK293F** | **HLJ** | 2.22 ± 0.33 | 6.18 ± 0.08 | **2.79 ± 0.03** | 1.41 ± 0.34 | 1.34 ± 0.28 | **9.50 ± 0.48** |
|  | **HEK293F** | **HL** | 4.65 ± 2.54 | 5.81 ± 2.61 | **1.25 ± 0.31** | 1.79 ± 0.51 | 0.73 ± 0.04 | **4.09 ± 0.17** |

As a conclusion, our work presents relevant results for the development of IgM as biopharmaceuticals, which requires new *in vitro* methods to produce biosimilars and to characterize the quality attributes of recombinant samples to meet the regulations of various drug and health authorities around the world. For antibodies, non-clinical studies must be performed to assess first the product qualities in terms of purity and homogeneity. Electrophoresis has been widely used so far but with the known limitations in detecting low protein amounts and the application of particular protocols necessary to characterize high molecular-weight biomolecules such as IgMs (Vorauer-Uhl et al., 2010). Our results showed the relevance of additional biophysical methods to precisely determine oligomeric distributions and the molecular weight of different IgM sample states. In particular, Mass Photometry appeared to be a reliable technique to assess rapidly the mass distribution with minimal sample amounts. The characterization of the similarity in known Fab- and Fc-associated functions such as binding to target antigen(s) and to Fc gamma receptors, FcRn or C1q are challenging, in particular in finding methods to demonstrate the binding of Igs to C1q. For now, the characterization of the latter case by ELISA have been widely used in pharmaceutical industry as a surrogate to complement dependent cytotoxicity assays in comparability studies for therapeutic Igs. Optical surface-based methods such as SPR and BLI have been proven to be alternatives with advantages like lower complexity in buffer preparations, lower hands-on manipulation, lower sample consumptions and to be faster, more in-depth and semi-automated Ig/C1q interaction analysis methods with higher precisions than the endpoint measurements from ELISA (Patel et al., 2015; Zhou et al., 2018; Jovic and Cymer, 2019). With our study, we demonstrated the suitability of the BLI-based assays for the interaction of recombinant IgM/C1q interactions and we believe that the developed methods and protocols can be easily applied to evaluate future IgM potential therapeutical preparations in combination with the conventional biological analysis.

## 5 Abbreviations

AUC: Analytical Ultra-Centrifugation
BLI: BioLayer Interferometry
CLR: Collagen-like region of C1q
ELISA: Enzyme-Linked Immunosorbent Assay
H/H chain: Heavy chain
L/L chain: Light chain
gC1q: globular head of C1q
Ig/Igs: Immunoglobuline/s
IgG/IgGs: Immunoglobuline/SG
IgM/IgMs: Immunoglobuline/sM
J/J chain: Joining chain
ELISA: Enzym-Linked Immunosorbent Assay
EM: Electron Microscopy
MP: Mass Photometry
PCR: Polymerase Chain Reaction
RIfS: Reflectometric Interference Spectroscopy
SEC: Size Exclusion Chromatography
SEC-MALLS: Multi-Angle-Light Scattering coupled to Size Exclusion Chromatography
SPR: Surface Plasmon Resonance
TEM: Transmitted Electron Microscopy
PAGE: PolyAcrylamide Gel Electrophoresis

## 6 Acknowledgments

cDNA coding for IgM617 and IgM012 chains were provided by Polymun Scientific Immunobiologische Forschung GmbH. This work used the Biophysical, AUC, EM and SPR/BLI platforms of the Grenoble Instruct-ERIC center (ISBG; UAR 3518 CNRS-CEA-UGA-EMBL) within the Grenoble Partnership for Structural Biology (PSB), supported by FRISBI (ANR-10-INBS-0005-02) and GRAL, financed within the University Grenoble Alpes graduate school (Ecoles Universitaires de Recherche) CBH-EUR-GS (ANR-17-EURE-0003). We thank Caroline Mas for her assistance at the Biophysical platform, Christine Ebel and Aline Le Roy at AUC platform. The EM facility is supported by the Auvergne Rhône-Alpes Region, the Fondation pour la Recherche Médicale (FRM), the Fonds FEDER and the GIS-Infrastructures en Biologie Santé et Agronomie (IBiSA). We thank Guy Schoehn, head of EM facility, for his support. IBS acknowledges integration into the Interdisciplinary Research Institute of Grenoble (IRIG, CEA). We would like to thank Véronique Rossi and Christine Gaboriaud from IBS for their expertise, helpful discussions and article proof reading.

## 8 Author Contributions

Conceptualization & supervision: Jean-Baptiste Reiser, Wai-Li Ling and Renate Kunert Funding acquisition: Nicole Thielens, Jean-Baptiste Reiser and Renate Kunert

Investigation and data: Julia Hennicke, Linda Schwaigerlehner and Renate Kunert (for IgM expression and purification in HEK293E and CHO DG44); Isabelle Bally, Anne Chouquet, Andrea Pinto and Jean-Baptiste Reiser (for IgM expression and purification in HEK293F); Anne Chouquet, Andrea Pinto and Jean-Baptiste Reiser (for Mass Photometry and BLI measurements), Andrea Pinto and Wai-Li Ling (for electron microscopy).

Expertise: Nicole Thielens (for C1r_2_C1s_2_ samples and SPR expertise).

Article writing: Jean-Baptiste Reiser, Wai-Li Ling, Andrea Pinto and Renate Kunert.

## References

Abdiche, Y., Malashock, D., Pinkerton, A., and Pons, J. (2008). Determining kinetics and affinities of protein interactions using a parallel real-time label-free biosensor, the Octet. Anal. Biochem. 377, 209–217. doi:10.1016/j.ab.2008.03.035.

Arlaud, G. J., Sim, R. B., Duplaa, A.-M., and Colomb, M. G. (1979). Differential elution of Clq, C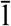r and C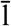s from human CT bound to immune aggregates. use in the rapid purification of C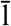 sub-components. Mol. Immunol. 16, 445–450. doi:10.1016/0161-5890(79)90069-5.

Arnold, J. N., Wormald, M. R., Suter, D. M., Radcliffe, C. M., Harvey, D. J., Dwek, R. A., et al. (2005). Human Serum IgM Glycosylation: IDENTIFICATION OF GLYCOFORMS THAT CAN BIND TO MANNAN-BINDING LECTIN*. J. Biol. Chem. 280, 29080–29087. doi:10.1074/jbc.M504528200.

Azuma, Y., Ishikawa, Y., Kawai, S., Tsunenari, T., Tsunoda, H., Igawa, T., et al. (2007). Recombinant Human Hexamer-Dominant IgM Monoclonal Antibody to Ganglioside GM3 for Treatment of Melanoma. Clin. Cancer Res. 13, 2745–2750. doi:10.1158/1078-0432.CCR-06-2919.

Bally, I., Inforzato, A., Dalonneau, F., Stravalaci, M., Bottazzi, B., Gaboriaud, C., et al. (2019). Interaction of C1q with pentraxin 3 and IgM revisited: mutational studies with recombinant C1q variants. Front. Immunol. 10, 461. doi:10.3389/fimmu.2019.00461.

Brautigam, C. A. (2015). Chapter Five - Calculations and Publication-Quality Illustrations for Analytical Ultracentrifugation Data. Methods Enzymol., 109–133. doi:10.1016/bs.mie.2015.05.001.

Calarese, D. A., Scanlan, C. N., Zwick, M. B., Deechongkit, S., Mimura, Y., Kunert, R., et al. (2003). Antibody Domain Exchange Is an Immunological Solution to Carbohydrate Cluster Recognition. Science 300, 2065–2071. doi:10.1126/science.1083182.

Cedzyński, M., Thielens, N. M., Mollnes, T. E., and Vorup-Jensen, T. (2019). Editorial: The Role of Complement in Health and Disease. Front. Immunol. 10, 1869. doi:10.3389/fimmu.2019.01869.

Chromikova, V., Mader, A., Hofbauer, S., Göbl, C., Madl, T., Gach, J. S., et al. (2015a). Introduction of germline residues improves the stability of anti-HIV mAb 2G12-IgM. Biochim. Biophys. Acta 1854, 1536–1544. doi:10.1016/j.bbapap.2015.02.018.

Chromikova, V., Mader, A., Steinfellner, W., and Kunert, R. (2015b). Evaluating the bottlenecks of recombinant IgM production in mammalian cells. Cytotechnology 67, 343–356. doi:10.1007/s10616-014-9693-4.

Collins, C., Tsui, F. W. L., and Shulman, M. J. (2002). Differential activation of human and guinea pig complement by pentameric and hexameric IgM. Eur. J. Immunol. 32, 1802–1810. doi:10.1002/1521-4141(200206)32:6<1802::AID-IMMU1802>3.0.CO;2-C.

Diebolder, C. A., Beurskens, F. J., de Jong, R. N., Koning, R. I., Strumane, K., Lindorfer, M. A., et al. (2014). Complement Is Activated by IgG Hexamers Assembled at the Cell Surface. Science 343, 1260–1263. doi:10.1126/science.1248943.

Doores, K. J., Fulton, Z., Huber, M., Wilson, I. A., and Burton, D. R. (2010). Antibody 2G12 Recognizes Di-Mannose Equivalently in Domain- and Nondomain-Exchanged Forms but Only Binds the HIV-1 Glycan Shield if Domain Exchanged. J. Virol. 84, 10690–10699. doi:10.1128/JVI.01110-10.

Duncan, A. R., and Winter, G. (1988). The binding site for C1q on IgG. Nature 332, 738. doi:10.1038/332738a0.

Eskeland, T., and Christensen, T. B. (1975). IgM Molecules with and without J Chain in Serum and after Purification, Studied by Ultra-centrifugation, Electrophoresis, and Electron Microscopy. Scand. J. Immunol. 4, 217–228. doi:10.1111/j.1365-3083.1975.tb02620.x.

Gasteiger, E., Hoogland, C., Gattiker, A., Duvaud, S., Wilkins, M. R., Appel, R. D., et al. (2005). “Protein Identification and Analysis Tools on the ExPASy Server,” in The Proteomics Protocols Handbook Springer Protocols Handbooks., ed. J. M. Walker (Totowa, NJ: Humana Press), 571–607. doi:10.1385/1-59259-890-0:571.

Gauglitz, G. (2020). Critical assessment of relevant methods in the field of biosensors with direct optical detection based on fibers and waveguides using plasmonic, resonance, and interference effects. Anal. Bioanal. Chem. 412, 3317–3349. doi:10.1007/s00216-020-02581-0.

Gilmour, J. E. M., Pittman, S., Nesbitt, R., and Scott, M. L. (2008). Effect of the presence or absence of J chain on expression of recombinant anti-Kell immunoglobulin M. Transfus. Med. 18, 167–174. doi:10.1111/j.1365-3148.2008.00853.x.

Gong, S., and Ruprecht, R. M. (2020). Immunoglobulin M: An Ancient Antiviral Weapon–Rediscovered. Front. Immunol. 11, 1943. doi:10.3389/fimmu.2020.01943.

Graille, M., Stura, E. A., Housden, N. G., Beckingham, J. A., Bottomley, S. P., Beale, D., et al. (2001). Complex between Peptostreptococcus magnus Protein L and a Human Antibody Reveals Structural Convergence in the Interaction Modes of Fab Binding Proteins. Structure 9, 679–687. doi:10.1016/S0969-2126(01)00630-X.

Hänel, C., and Gauglitz, G. (2002). Comparison of reflectometric interference spectroscopy with other instruments for label-free optical detection. Anal. Bioanal. Chem. 372, 91–100. doi:10.1007/s00216-001-1197-3.

Harboe, M., Thorgersen, E. B., and Mollnes, T. E. (2011). Advances in assay of complement function and activation. Adv. Drug Deliv. Rev. 63, 976–987. doi:10.1016/j.addr.2011.05.010.

Hennicke, J., Lastin, A. M., Reinhart, D., Grünwald-Gruber, C., Altmann, F., and Kunert, R. (2017). Glycan profile of CHO derived IgM purified by highly efficient single step affinity chromatography. Anal. Biochem. 539, 162–166. doi:10.1016/j.ab.2017.10.020.

Hennicke, J., Reinhart, D., Altmann, F., and Kunert, R. (2019). Impact of temperature and pH on recombinant human IgM quality attributes and productivity. New Biotechnol. 50, 20–26. doi:10.1016/j.nbt.2019.01.001.

Hennicke, J., Schwaigerlehner, L., Grünwald-Gruber, C., Bally, I., Ling, W. L., Thielens, N., et al. (2020). Transient pentameric IgM fulfill biological function—Effect of expression host and transfection on IgM properties. PLOS ONE 15, e0229992. doi:10.1371/journal.pone.0229992.

Hiramoto, E., Tsutsumi, A., Suzuki, R., Matsuoka, S., Arai, S., Kikkawa, M., et al. (2018). The IgM pentamer is an asymmetric pentagon with an open groove that binds the AIM protein. Sci. Adv. 4, eaau1199. doi:10.1126/sciadv.aau1199.

Idusogie, E. E., Presta, L. G., Gazzano-Santoro, H., Totpal, K., Wong, P. Y., Ultsch, M., et al. (2000). Mapping of the C1q Binding Site on Rituxan, a Chimeric Antibody with a Human IgG1 Fc. J. Immunol. 164, 4178–4184. doi:10.4049/jimmunol.164.8.4178.

Jones, K., Savulescu, A. F., Brombacher, F., and Hadebe, S. (2020). Immunoglobulin M in Health and Diseases: How Far Have We Come and What Next? Front. Immunol. 11. doi:10.3389/fimmu.2020.595535.

Jovic, M., and Cymer, F. (2019). Qualification of a surface plasmon resonance assay to determine binding of IgG-type antibodies to complement component C1q. Biologicals 61, 76–79. doi:10.1016/j.biologicals.2019.08.004.

Keyt, B. A., Baliga, R., Sinclair, A. M., Carroll, S. F., and Peterson, M. S. (2020). Structure, Function, and Therapeutic Use of IgM Antibodies. Antibodies 9, 53. doi:10.3390/antib9040053.

Kishore, U., and Reid, K. B. M. (2000). C1q: Structure, function, and receptors. Immunopharmacology 49, 159–170. doi:10.1016/S0162-3109(00)80301-X.

Kumar, N., Arthur, C. P., Ciferri, C., and Matsumoto, M. L. (2021). Structure of the human secretory immunoglobulin M core. Structure, S0969212621000022. doi:10.1016/j.str.2021.01.002.

Le Roy, A., Nury, H., Wiseman, B., Sarwan, J., Jault, J.-M., and Ebel, C. (2013). “Sedimentation Velocity Analytical Ultracentrifugation in Hydrogenated and Deuterated Solvents for the Characterization of Membrane Proteins,” in Membrane Biogenesis: Methods and Protocols Methods in Molecular Biology., eds. D. Rapaport and J. M. Herrmann (Totowa, NJ: Humana Press), 219–251. doi:10.1007/978-1-62703-487-6_15.

Le Roy, A., Wang, K., Schaack, B., Schuck, P., Breyton, C., and Ebel, C. (2015). “Chapter Twelve - AUC and Small-Angle Scattering for Membrane Proteins,” in Methods in Enzymology Analytical Ultracentrifugation., ed. J. L. Cole (Academic Press), 257–286. doi:10.1016/bs.mie.2015.06.010.

Li, Y., Wang, G., Li, N., Wang, Y., Zhu, Q., Chu, H., et al. (2020). Structural insights into immunoglobulin M. Science 367, 1014–1017. doi:10.1126/science.aaz5425.

Mader, A., Chromikova, V., and Kunert, R. (2013). Recombinant IgM expression in mammalian cells: A target protein challenging biotechnological production. Adv. Biosci. Biotechnol. 04, 38–43. doi:10.4236/abb.2013.44A006.

Moore, G. L., Chen, H., Karki, S., and A, G. (2010). Engineered Fc variant antibodies with enhanced ability to recruit complement and mediate effector functions. mAbs 2, 181–189. doi:10.4161/mabs.2.2.11158.

Patel, R., Neill, A., Liu, H., and Andrien, B. (2015). IgG subclass specificity to C1q determined by surface plasmon resonance using Protein L capture technique. Anal. Biochem. 479, 15–17. doi:10.1016/j.ab.2015.03.012.

Randall, T. D., King, L. B., and Corley, R. B. (1990). The biological effects of IgM hexamer formation. Eur. J. Immunol. 20, 1971–1979. doi:10.1002/eji.1830200915.

Randall, T. D., Parkhouse, R. M., and Corley, R. B. (1992). J chain synthesis and secretion of hexameric IgM is differentially regulated by lipopolysaccharide and interleukin 5. Proc. Natl. Acad. Sci. U. S. A. 89, 962–966. doi:10.1073/pnas.89.3.962.

Ricklin, D., Hajishengallis, G., Yang, K., and Lambris, J. D. (2010). Complement - a key system for immune surveillance and homeostasis. Nat. Immunol. 11, 785–797. doi:10.1038/ni.1923.

Salvay, A. G., Santamaria, M., le Maire, M., and Ebel, C. (2008). Analytical Ultracentrifugation Sedimentation Velocity for the Characterization of Detergent-Solubilized Membrane Proteins Ca++-ATPase and ExbB. J. Biol. Phys. 33, 399. doi:10.1007/s10867-008-9058-3.

Schuck, P. (2000). Size-Distribution Analysis of Macromolecules by Sedimentation Velocity Ultracentrifugation and Lamm Equation Modeling. Biophys. J. 78, 1606–1619. doi:10.1016/S0006-3495(00)76713-0.

Sharp, T. H., Boyle, A. L., Diebolder, C. A., Kros, A., Koster, A. J., and Gros, P. (2019). Insights into IgM-mediated complement activation based on in situ structures of IgM-C1-C4b. Proc. Natl. Acad. Sci., 201901841. doi:10.1073/pnas.1901841116.

Sonn-Segev, A., Belacic, K., Bodrug, T., Young, G., VanderLinden, R. T., Schulman, B. A., et al. (2020). Quantifying the heterogeneity of macromolecular machines by mass photometry. Nat. Commun. 11, 1772. doi:10.1038/s41467-020-15642-w.

Sultana, A., and Lee, J. E. (2015). Measuring Protein-Protein and Protein-Nucleic Acid Interactions by Biolayer Interferometry. Curr. Protoc. Protein Sci. 79, 19.25.1-19.25.26. doi:10.1002/0471140864.ps1925s79.

Ugurlar, D., Howes, S. C., de Kreuk, B.-J., Koning, R. I., de Jong, R. N., Beurskens, F. J., et al. (2018). Structures of C1-IgG1 provide insights into how danger pattern recognition activates complement. Science 359, 794–797. doi:10.1126/science.aao4988.

Vorauer-Uhl, K., Wallner, J., Lhota, G., Katinger, H., and Kunert, R. (2010). IgM characterization directly performed in crude culture supernatants by a new simple electrophoretic method. J. Immunol. Methods 359, 21–27. doi:10.1016/j.jim.2010.05.003.

Wiersma, E. J., Collins, C., Fazel, S., and Shulman, M. J. (1998). Structural and Functional Analysis of J Chain-Deficient IgM. J. Immunol. 160, 5979–5989.

Wolbank, S., Kunert, R., Stiegler, G., and Katinger, H. (2003). Characterization of Human Class-Switched Polymeric (Immunoglobulin M [IgM] and IgA) Anti-Human Immunodeficiency Virus Type 1 Antibodies 2F5 and 2G12. J. Virol. 77, 4095–4103. doi:10.1128/JVI.77.7.4095-4103.2003.

Zhao, H., Ghirlando, R., Alfonso, C., Arisaka, F., Attali, I., Bain, D. L., et al. (2015). A Multilaboratory Comparison of Calibration Accuracy and the Performance of External References in Analytical Ultracentrifugation. PLoS ONE 10, e0126420. doi:10.1371/journal.pone.0126420.

Zhou, W., Lin, S., Chen, R., Liu, J., and Li, Y. (2018). Characterization of antibody-C1q interactions by Biolayer Interferometry. Anal. Biochem. 549, 143–148. doi:10.1016/j.ab.2018.03.022.

