## Supplementary figures for "Biophysical characterization of the oligomeric states of recombinant Immunoglobulins type-M and their C1q binding kinetics by Biolayer Interferometry"

### Supplementary Figure S1

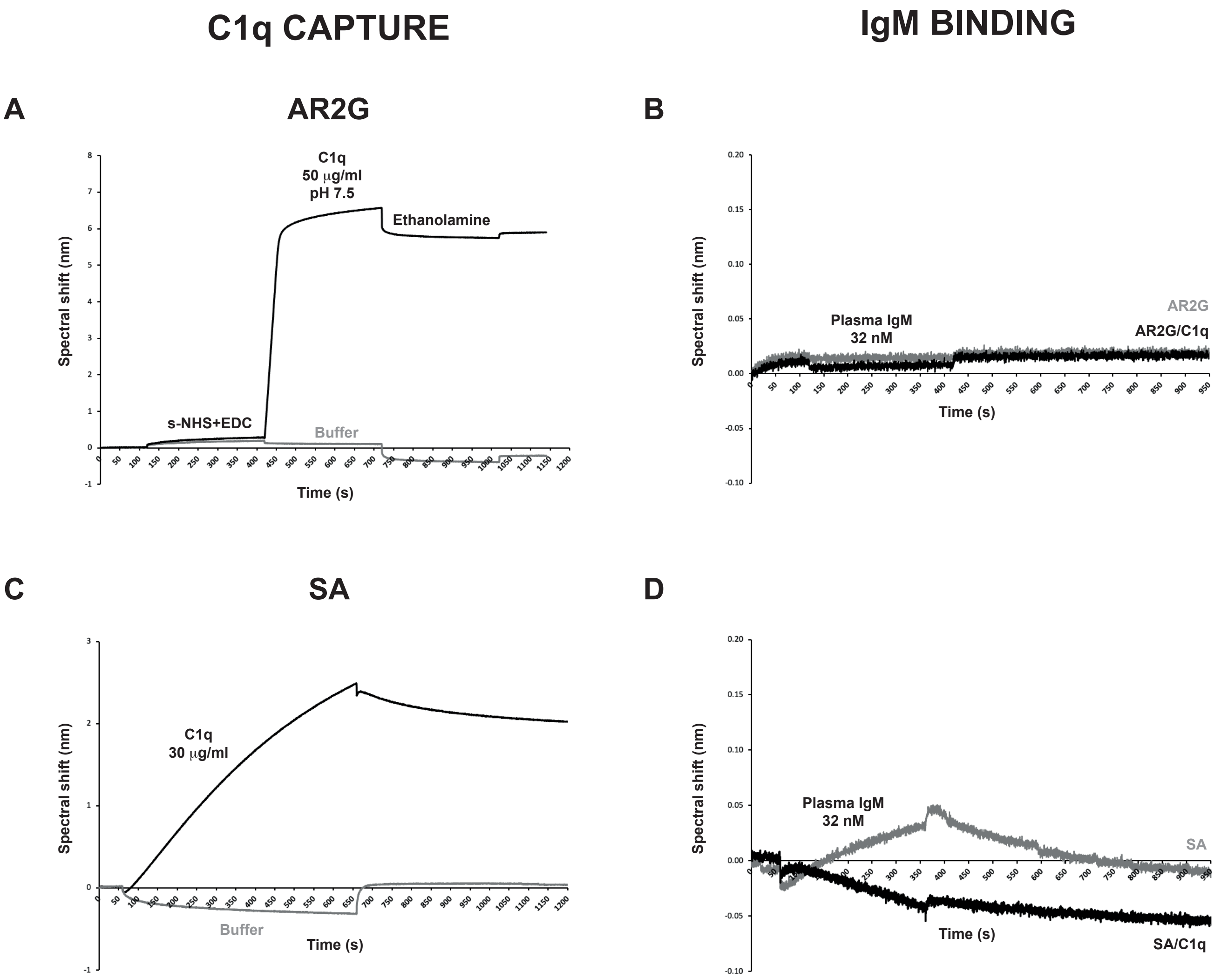


**Supplementary Figure S1.** Exemples of C1q captures on (**A**) AR2G and (**C**) SA biosensors and (**B and D**) of plasma IgM binding at 32 nM to functionalyzed (in black) and reference (in grey) biosensors.

### Supplementary Figure S2

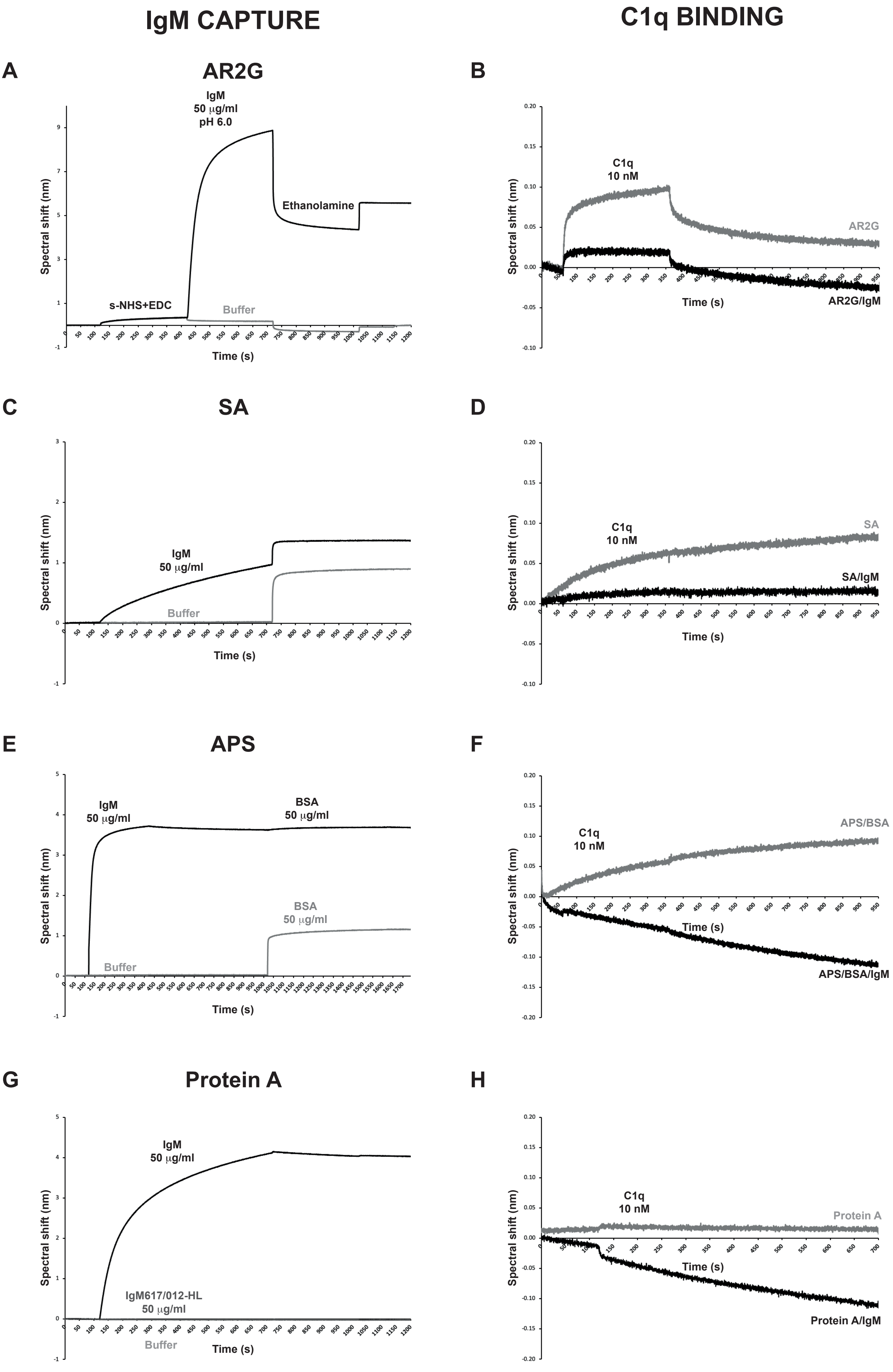

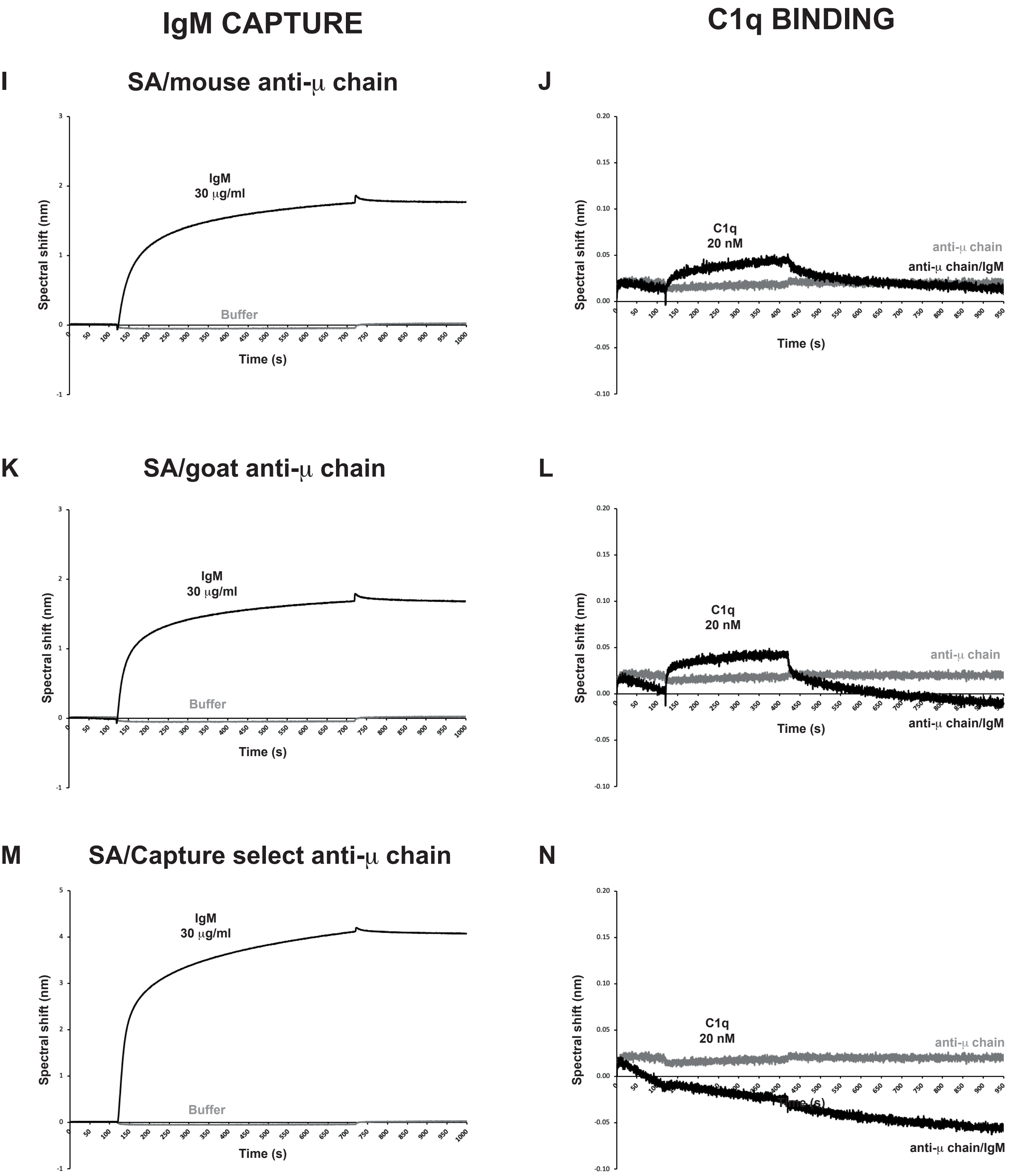


**Supplementary Figure S2.** Examples of plasma IgM captures on (**A**) AR2G, (**C**) SA, (**E**) APS, (**G**) Protein A, (**I, K, M**) anti-μ chain biosensors and (**B, D, F, H, J, L, N**) of plasma C1q binding at 10 or 20 nM to functionalyzed (in black) and reference (in grey) biosensors.

### Supplementary Figure S3

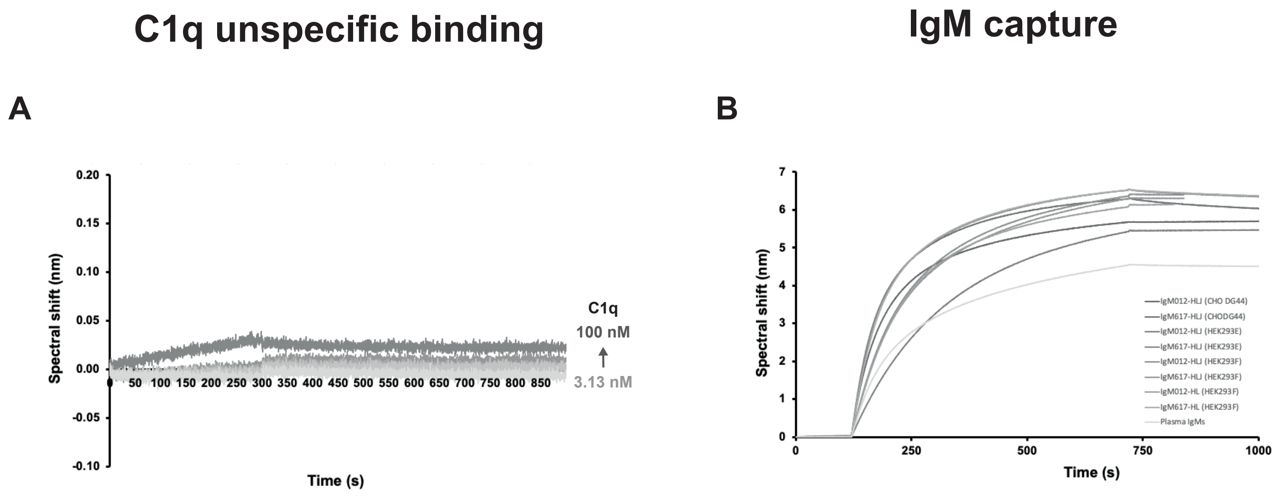


**Supplementary Figure S3**. (**A**) C1q unspecific binding on Protein L biosensor at different concentrations from 3.13 nM to 100 nM. (**B**) Capture of IgM samples onto Protein L biosensors (IgM concentration = 30 µg/ml).

### Supplementary Figure S4

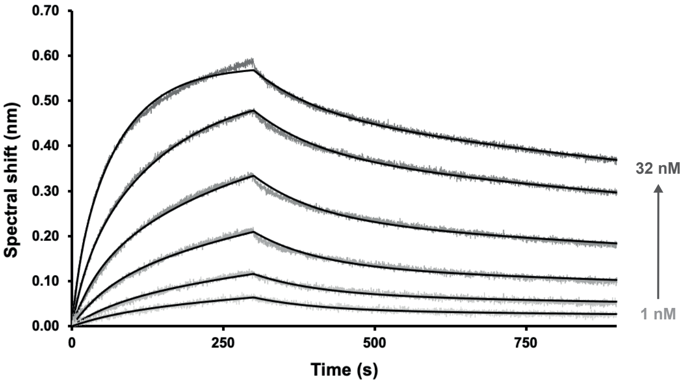


| $k_{a1}$ | $k_{d1}$ | **$K_{D1}$** | $k_{a2}$ | $k_{d2}$ | **$K_{D2}$** |
|---|---|---|---|---|---|
| $10^5$/Ms | $10^{-4}$/s | **$10^{-9}$ M** | $10^6$/Ms | $10^{-3}$/s | **$10^{-9}$ M** |
| 4.06 ± 0.01 | 4.02 ± 0.03 | **10.00 ± 0.07** | 2.14 ± 0.02 | 9.28 ± 0.01 | **4.33 ± 0.01** |

**Supplementary Figure S4.** Kinetics analysis of the interaction between C1q from plasma and recombinant $C1r_2C1s_2$. C1q was immobilized on AR2G biosensor (supplementary Figure S1). The tetramer was expressed in mammalian expression (Bally et al., 2019). The functionalized biosensors were dipped in wells containing $C1r_2C1s_2$ at different concentrations (1, 2, 4, 8, 16, 32 nM). The binding signals (grey-scaled sensorgrams) were obtained by subtracting the signals from empty AR2G biosensor and from zero-concentration samples. Fitted curved are depicted as black lines and were obtained by global fitting using a 2:1 heterogenous ligand model. Kinetics parameters and affinities are reported in the table.
